# Defining the synaptic mechanisms that tune CA3-CA1 reactivation during sharp-wave ripples

**DOI:** 10.1101/164699

**Authors:** Paola Malerba, Matt W. Jones, Maxim A. Bazhenov

## Abstract

During non-REM sleep, memory consolidation is driven by a dialogue between hippocampus and cortex involving the reactivation of specific neural activity sequences (‘replay’). In the hippocampus, replay occurs during sharp-wave ripples (SWRs), short bouts of excitatory activity in area CA3 which induce high frequency oscillations in the inhibitory interneurons of area CA1. Despite growing evidence for the functional importance of replay, its neural mechanisms remain poorly understood. Here, we develop a novel theoretical model of hippocampal spiking during SWRs. In our model, noise-induced activation of CA3 pyramidal cells triggered an excitatory cascade capable of inducing local ripple events in CA1. Ripples occurred stochastically, with Schaffer Collaterals driving their coordination, so that localized sharp waves in CA3 produced consistently localized CA1 ripples. In agreement with experimental data, the majority of pyramidal cells in the model showed low reactivation probabilities across SWRs. We found, however, that a subpopulation of pyramidal cells had high reactivation probabilities, which derived from fine-tuning of the network connectivity. In particular, the excitatory inputs along synaptic pathway(s) converging onto cells and cell pairs controlled emergent single cell and cell pair reactivation, with inhibitory inputs and intrinsic cell excitability playing differential roles in CA3 vs. CA1. Our model predicts (1) that the hippocampal network structure driving the emergence of SWR is also able to generate and modulate reactivation, (2) inhibition plays a particularly prominent role in CA3 reactivation and (3) CA1 sequence reactivation is reliant on CA3-CA1 interactions rather than an intrinsic CA1 process.

## New and Noteworthy

We develop a new biophysical model of hippocampal sharp-wave ripple activity to study determinants of pyramidal cell reactivation during sleep. The majority of pyramidal cells revealed low reactivation probabilities arising from stationary network properties. Highly reactivating subpopulations were driven by cell-specific synaptic inputs, with CA3 excitatory recurrents and CA3-CA1 Schaffer Collaterals interacting to promote reliable CA1 sequence reactivation. The model recapitulates experimental data and highlights mechanisms by which experience-dependent inputs can tune memory consolidation.

## Introduction

Memories acquired during wakefulness continue to evolve during subsequent sleep. Sleep seems an optimal brain state for this memory consolidation: the brain is dissociated from external inputs and internal processing can be supported by sleep stage-dependent patterns of network activity, driven largely by periodic shifts in neuromodulatory tone (Krishnan et al. 2016; Lee and Dan 2012). During non-REM sleep, hippocampal networks show sharp-wave ripples (SWR), short bouts of synchronized population activity (50-100ms) initiated in the CA3 area of the hippocampus with strong excitatory firing that reaches area CA1, driving fast spiking interneurons to rhythmically organize a small fraction of local pyramidal cells spiking (Buzsaki 2015; Buzsaki et al. 1983; Mizuseki et al. 2012). In CA1, local field potential (LFP) high frequency oscillations (above 150Hz in the rat, about 100Hz in humans) occur in the pyramidal layer (the ripple), while in *stratum radiatum* a strong deflection marks the effects of Schafferal Collateral input to the pyramidal cells (the sharp wave). Hippocampal SWR can be locked to cortical slow oscillations (SO) and troughs of thalamocortical spindles, in a coordination of activity across brain regions and time scales (Molle and Born 2011; Sirota and Buzsaki 2005), which is thought to orchestrate a hippocampal-neocortical dialogue mediating memory consolidation. In fact, it is known that the number of SWR correlates with memory performance after sleep (Eschenko et al. 2008; Ramadan et al. 2009), suppressing SWR compromises memory consolidation (Ego-Stengel and Wilson 2010; Girardeau et al. 2009) and increased SO power and coordination of SO and other sleep rhythms augments memory (Marshall et al. 2006).

Reactivation of specific neural activity patterns – replay – during slow wave sleep has been observed in both hippocampus and neocortex (Hoffman and McNaughton 2002; Ji and Wilson 2007; Schwindel and McNaughton 2011; Skaggs and McNaughton 1996; Sutherland and McNaughton 2000; Wilson and McNaughton 1994), and coincides with SWR (Foster and Wilson 2006; Wu and Foster 2014). Together, these facts led to the hypothesis that coordinated sequence reactivation during precisely timed oscillations can recruit synaptic plasticity, leading to memory consolidation across brain structures (Diekelmann and Born 2010). Within this hypothesis, understanding how replay happens in hippocampal SWR is crucial to explain sleep dependent memory consolidation. More specifically, it is important to understand how hippocampal replay depends on awake activity and in particular on the hippocampal synaptic connections which are known to change rapidly during learning. It remains unknown whether changing only a small fraction of CA1-CA3 synapses during learning can be sufficient to provide reliable reactivation of these synapses and neurons they connect during sleep and without changing the overall network dynamics.

To date, the mechanisms underlying sequence reactivation during SWRs are not known. One hypothesis is that input-driven partial reactivation in CA3 can perform pattern completion, which in turns drives a refined CA1 spike reactivation. Sequential cell reactivation has primarily been demonstrated in CA1 pyramidal cells during SWRs, but fewer data are available on CA3 pyramidal cell replay (Carr et al. 2012). Hence it is not clear if reactivation of CA1 sequences is shaped by intrinsic CA1 mechanisms or depends on CA3 input, and possible CA3 sequence reactivation. To address this question, we introduced a computational model of CA3-CA1 SWR activity, and we studied SWR-mediated reactivation in both regions. Our model revealed large CA3 excitatory events (the sharp waves), occurring spontaneously, and driving rhythmic spiking in CA1 interneurons (the ripples). This model was applied to study how the spontaneous SWR dynamics in the network influences the distribution of cell activation within CA3 and CA1 pyramidal cells populations, and how specific inputs that can be modified during learning can act on selected cells within these populations to promote their reactivation. It predicts that the same network structure that drives SWR is also able to generate and modulate reactivation, and that inhibition and intrinsic cell excitability play a role in CA3 reactivation but not CA1. Finally, our model proposes that CA3-CA1 synaptic architecture has to be changed as a unified system to promote reliable CA1 replay.

## Results

### Computational model of spontaneous, localized SPW-R activity

In our previous work (Malerba et al. 2016), we introduced a model of CA1 ripples in which local field potential (LFP) oscillations were due to a transient in the system dynamics, as opposed to a stable oscillatory state. Transients were imposed by fast firing of the basket cells initially synchronized by common input current (representing CA3 excitation); cell population heterogeneity led to loss of coordination over time and ripple termination. Here, we build on that work to introduce a model of CA3-CA1 SWRs in which CA3 activity emerges spontaneously and triggers stochastic activation of SWR events in CA3 and CA1, with the termination of ripples driven by the same de-coordination mechanism. In the new model, different subsets of cells participated in different SWR events as observed experimentally (Patel et al. 2013).

The model of CA3-CA1 SWR activity is based on synaptically coupled populations of pyramidal cells and basket cells (Figure 1A). It includes highly recurrent strong excitatory AMPA receptor-mediated connections between CA3 pyramidal cells, and weak and sparse recurrent excitatory connections within CA1 pyramidal cells (Shepherd 2004). CA3 pyramidal cells projected excitatory connections to CA1 cells, representing the Schaffer Collaterals. The CA3 network and its projections to CA1 had stochastic densities and strengths within a radius of about a third of the target network (Figure 1B shows in a matrix the presence of synaptic connections), which is consistent with analysis of CA3 pyramidal cells arborization (Li et al. 1994). Importantly, each neuron received an independent noise current which drove occasional irregular spiking, and a baseline constant drive which was selected from a Gaussian distribution. Details of the computational model rationale and equations are reported in Materials and Methods.

**Figure 1.**
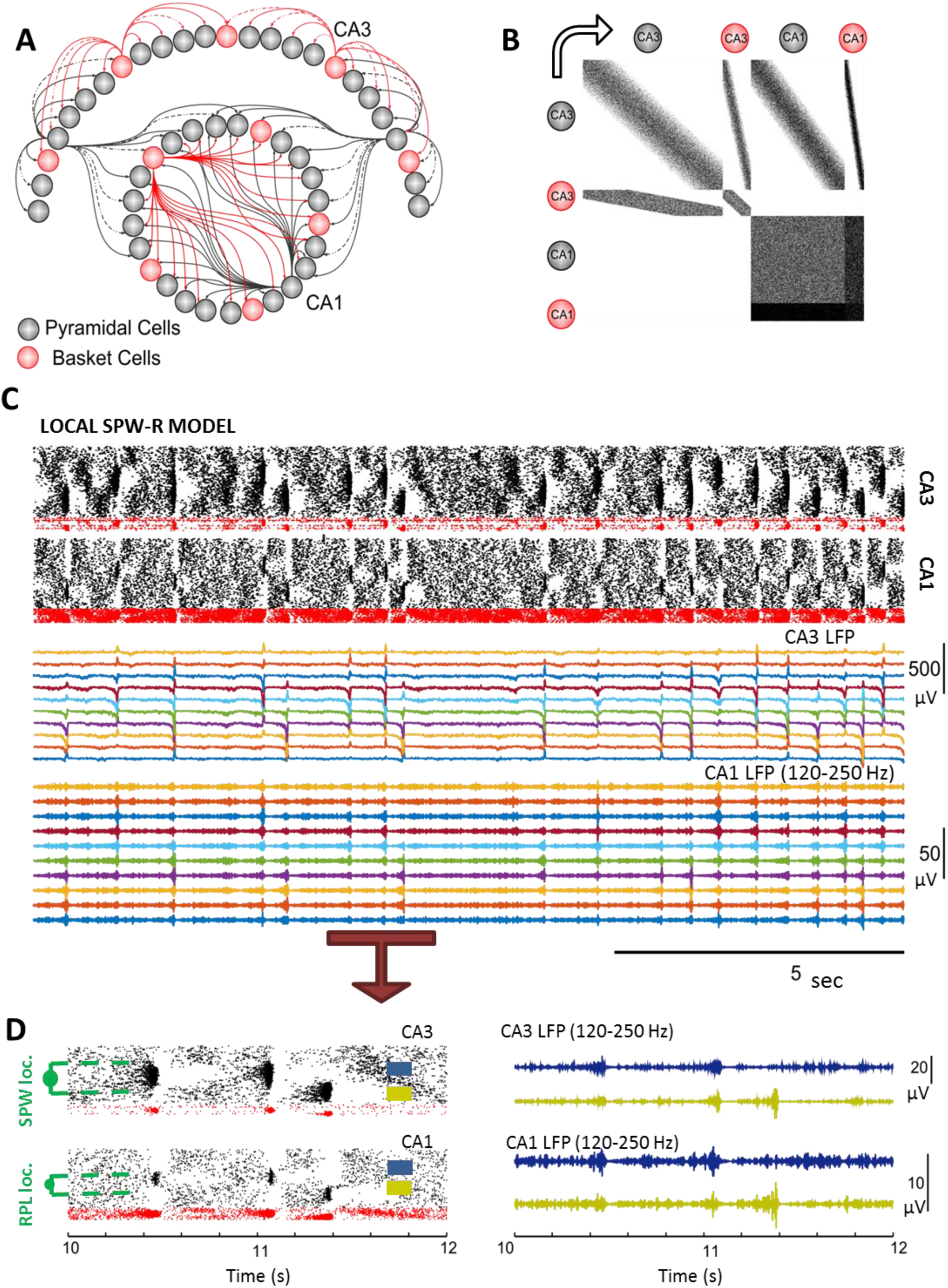
Computational Model of emergent, localized SPW-R activity. ***A.*** *Schematic representation of the model, showing the two main regions (CA3 and CA1) and the cell types considered (pyramidal cells and basket cells). Note that CA3 projects to CA1, but not vice-versa.* ***B.*** *Matrix representation of presence of synaptic connections in the model. Synaptic weights are not shown, hence darker tone is only indicative of local higher density of connections. Note that CA3 pyramidal cells connect to all cell types in the network.* ***C.*** *Example of SPW-R activity in the network. Top 2 plots*: *raster plots of cell spikes (CA3 above, CA1 below). Dots mark spikes in time of pyramidal cells (black) and interneurons (red). In CA3 sharp waves happen at different locations and show different propagation patterns in time. CA1 spiking is organized by the sharp waves in CA3, and ripples are visible as small sharp stripes of dense spiking in CA1 pyramidal cells. Bottom two plots*: *Local Field Potential (LFP) of the model, computed as the average of the total incoming synaptic currents across a group of pyramidal cells. Note that in CA3 we show the wide-band signal, to highlight the sharp transition occurring in the synaptic currents when a sharp wave is present in the CA3 network. At corresponding times (and locations) in the LFP of CA1 (filtered in ripple range) one can see the high frequency activity captured by the LFP signal.* ***D.*** *Zoomed-in raster plot of spiking activity in CA3 and CA1, the time window is indicated by the arrow in C. SWR activity is localized within the two regions. For each sharp wave (SPW) and each ripple (RPL) a center (or location) can be defined as the medium index among the pyramidal cells which spike during the event. The LFPs shown on the right refer to groups of cells slightly apart in the network, shown as colored rectangles in the rastergram. Note that some SWR can be seen in the LFP traces at both locations (near 11s) while others are only visible in one of the traces (about 400ms later).*

As shown in Figure 1C, the network spontaneously organized into bouts of CA3 pyramidal cell spiking, which drove spiking in CA1. In CA1, interneurons organized their firing into high frequency oscillations, and a few pyramidal cells spiked within windows of opportunity left at the troughs of the lateral synaptic inhibition oscillations, thus forming a SWR event. As is typical in real data (Davidson et al. 2009), SWRs occurred in temporal clusters punctuated by long pauses. A representation of the LFP obtained by averaging the synaptic currents impinging on subsets of pyramidal cells showed that SWR events in CA3 and CA1 were localized, and the location of the SWRs within the network changed in time. This is consistent with experimental findings showing that ripple events can be localized in space (Patel et al. 2013; Ylinen et al. 1995) and that CA3 pyramidal cells are known to be very active during SWR, but do not spike phase-locked to CA1 ripples (Atherton et al. 2015; Sullivan et al. 2011).

Figure 1D shows a zoomed-in version of the SWR spiking activity and LFPs in sub-regions of CA3 and CA1 which were connected by Schaffer Collaterals. Although sharp waves are typically experimentally measured in CA1 *stratum radiatum*, in the following we refer to sharp waves as the bouts of excitatory activity in CA3 which lead to ripples in CA1. The general activity of the model was consistent with known properties of SWRs: ripple frequency was 174±21.3 Hz, ripple durations were 54±27 ms, sharp wave durations were 126±23ms and the inter-event pauses (time durations between two successive sharp waves in CA3 or ripples in CA1) showed approximately exponential distributions, which fitted to exponential functions with rates of 1.08 Hz in CA3 and 1.2Hz in CA1 (Buzsaki 2015). Within this model, we studied the spontaneous activation of CA3 and CA1 pyramidal cells across multiple ripples, in relation to their synaptic properties.

### Activation during SWR is a basic sampling process for low activation cells, but not for highly activating cells

We started our analysis by quantifying cell activation during ripples using an “R-activation score”: the percent of all ripples in which a given pyramidal cell spiked at least once (Fig 2 A-B). On average across 20, 100s simulations, we found that CA3 pyramidal cells activated more often than CA1 cells, likely because sharp waves involve about 30% of all cells in the CA3 network while ripples only involve about 15% of all CA1 pyramidal cells (consistent with data (Csicsvari et al. 1999a; b; Sullivan et al. 2011), not shown). Since in our model SWR are localized (involve only a portion of the network) and erratic (they happen in different locations over time), we reasoned that R-activation of any cell could be determined by how often a SWR happened in the vicinity of that cell. If that was the case, than the R-activation of any cell in the network would be the result of a sampling process, where each SWR event is one independent sample of a few cells within the network (30% and 15%, for CA3 and CA1 respectively), and cells are present in any sample according to a stationary (fixed in time) probability.

**Figure 2.**
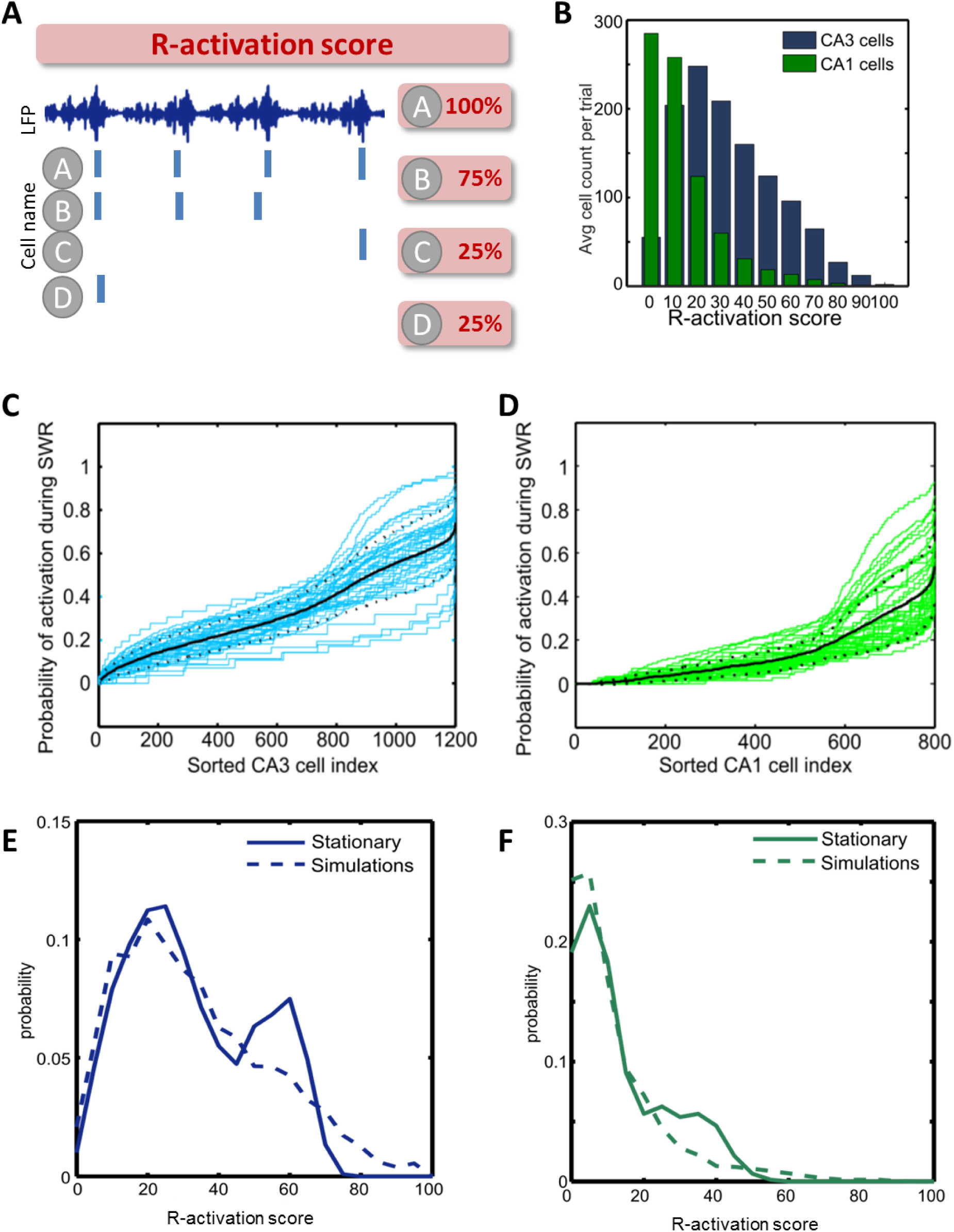
R-activation in CA3 and CA1 is stationary for low score cells and dynamic for high score cells. ***A.*** *Drawing shows the definition of R-activation score for a pyramidal cell. The total number of SWR in which a cell spikes during a 100 second simulation is found, and compared to the total number of SWR in the simulation. This fraction is expressed as a percent as the R-activation score of the cell.* ***B***. *Distribution of R-activation scores of CA3 and CA1 pyramidal cells computed across many simulations and reported as average count in any given single trial. CA1 pyramidal cells show a peak for 0% R-activation, and the mean R-activation scores for CA3 pyramidal cells is higher than CA1 pyramidal cells.* ***C.*** *Curves (one per simulation) mark the probability distribution of pyramidal cells in CA3 to be spiking in any given SWR. Cells were sorted by increasing probabilities. The average probability curve (across all curves for each sorted cell index) is marked by a black solid line, while dotted black lines represent the standard deviation around the mean.* ***D.*** *Same as panel C for CA1 pyramidal cells.* ***E.*** *Distributions of R-activation scores in CA3 pyramidal cells in a stationary sorting algorithm (thick line) and in the model (dotted line). Note that the stationary process and the model share the low-reactivation peak probability, but have different trends for high-reactivations: the stationary choice peaks at 60% and quickly decays to zero for higher scores, while the model does not show peaks at high R-activation values (only the low R-activation peak is present) and has a larger amount of very high R-activations.* ***F.*** *Same as panel E for CA1 pyramidal cells: again stationary choice and computational model share the low R-activation peak and the model does not show a peak for intermediate levels of R-activation.*

We tested this idea by finding an average stationary probability of spiking during SWR for CA3 and CA1 cells (Fig 2 C-D), and running a simple sampling process over such probabilities (see Materials and Methods). We then compared the distribution of R-activation scores found in the sampling process and in the simulations (Fig 2 E-F) and found that for low R-activations, the biophysical model and the sampling process showed very similar profiles, peaking at 20% for CA3 and 10% for CA1. At high R-activations the sampling process showed a second peak (60% for CA3 and 35% for CA1) which was not present in the biophysical model. Hence, the distributions of R-activation scores in our model were equivalent to a simple sampling process for cells with low reactivation scores (the majority in both networks), but not for cells at higher scores. In fact, the lack of a second peak was replaced by the presence of a tail in the distributions (which is not present in the sampling distributions). These tails showed that in both CA3 and CA1 there were few cells with high R-activation scores, and that their presence could not be explained as a result of a simple sampling process. This is particularly interesting because data show that during ripples CA1 cells can be separated in high-firing and low-firing according to how often they activate across ripples (where the high firing cells code for recently formed memories) (Grosmark and Buzsaki 2016; Mizuseki et al. 2012), and our model suggests that low and high firing cells emerge from two superimposed mechanisms: a basic sampling process for the low R-activating cells and a cell-specific selection process for the high R-activating cells. This selection process could identify specific network and cell characteristics that can be changed during learning to promote the reactivation of recently acquired memories in the hippocampus.

### The influence of input on pyramidal cell R-activation

We set out to understand which synaptic and intrinsic inputs could impact the reactivation of cells. We quantified inputs differently in CA3 and CA1 (formulae on Figure 3), but in all cases considered the role of excitatory synaptic input, inhibitory synaptic input and intrinsic excitability of the cell, and estimated the relative role of the different inputs on R-activation by multivariate linear regression analysis (tables available in supporting information)(Figure 3). Details of how the input quantifiers were defined are reported in Appendix.

**Figure 3.**
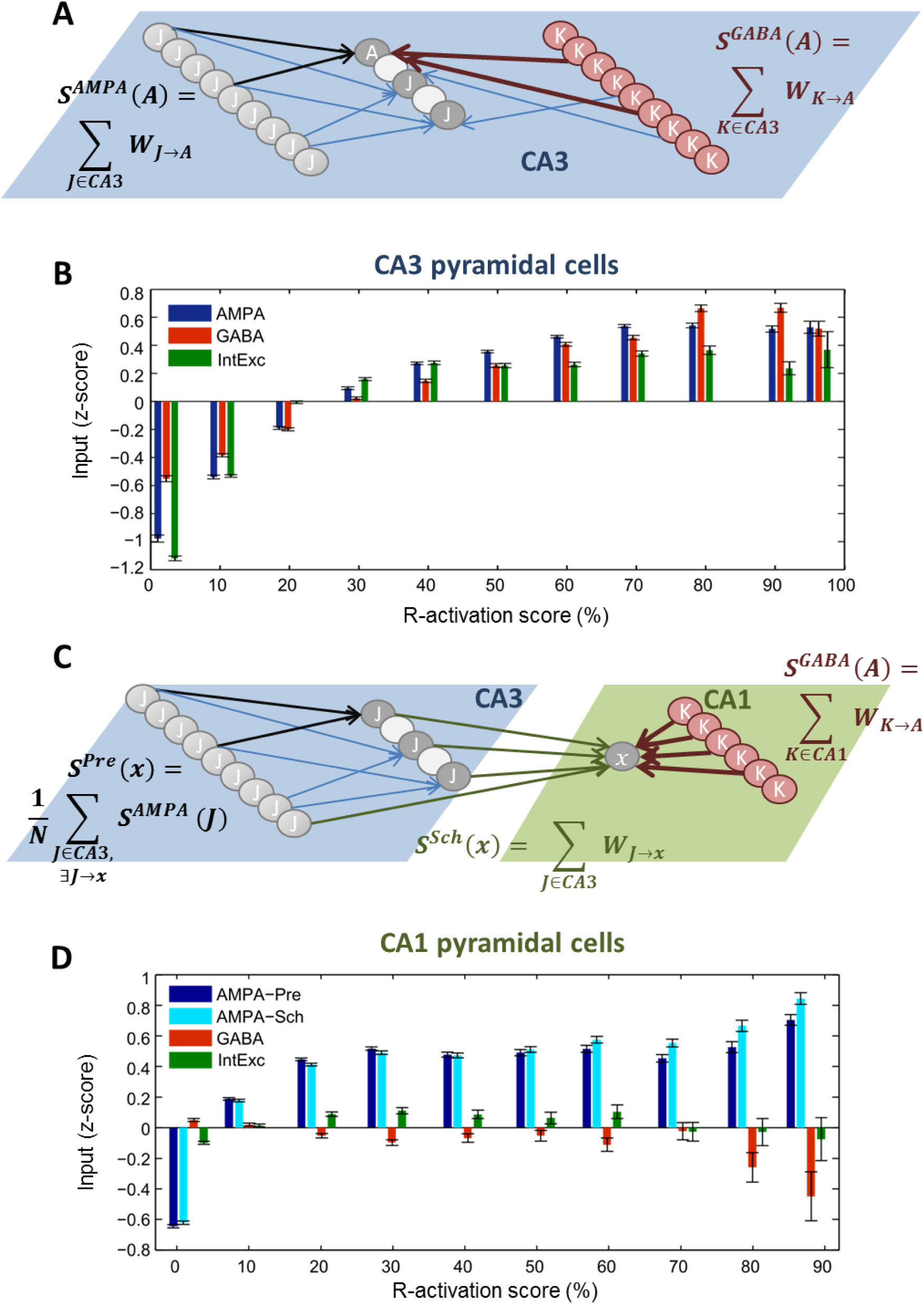
In CA3 and CA1, pyramidal cell R-activation scores increase with excitatory input. ***A.*** *Drawing of the synaptic inputs analyzed for CA3 pyramidal cells. S*^*AMPA*^ *is the sum of all incoming AMPA connections (in black) to a CA3 pyramidal cell (labeled A) from other CA3 pyramidal cells (in gray). S*^*GABA*^ *is the sum of all incoming GABA connections (in red) to a CA3 pyramidal cell (A) from CA3 interneurons (in red).* ***B.*** *R-activation of CA3 pyramidal cells is related to strength of synaptic inputs and intrinsic excitability. The bar plot shows on the x –coordinate the average R-activation score of CA3 pyramidal cells belonging to the same score group (±5%) and on the y-coordinate the level of each different input (S*^*AMPA*^, *S*^*GABA*^ *or Intrinsic Excitability) for cells within that score group. Error bars mark the standard error of the mean. Before grouping cells by their R-activation scores, each input was z-scored to enable comparisons of their respective trends. Therefore, a negative y-coordinate does not reflect a negative input, but an input below the average across the whole network.* ***C.*** *Drawing of the synaptic inputs analyzed for CA1 pyramidal cells. S*^*Sch*^ *(from CA3 to CA1, in green) is the sum of incoming AMPA synaptic weights from CA3 cells onto CA1 pyramidal cells. S*^*Pre*^ *(in black) for a given CA1 pyramidal cell (labeled x) finds all cells in CA3 that projects to x and their AMPA input (S*^*AMPA*^, *described in* ***A****). The average value of all these S*^*AMPA*^ *inputs is S*^*Sch*^, *representing the how much excitatory drive the cells in CA3 which project to x in CA1 are receiving. S*^*GABA*^ *is the sum of all incoming GABA connections (in red) to a CA1 pyramidal cell (x) from CA1 interneurons (in red).* ***D.*** *R-activation of CA1 pyramidal cells is related to synaptic excitatory input. The bar plot shows on the x –coordinate the average R-activation score of CA1 pyramidal cells belonging to the same score group (±5%) and on the y-coordinate the level of each different input (S*^*Pre*^, *S*^*Sch*^ *or Intrinsic Excitability) for cells within that score group. Inputs were z-scored before grouping the cells by score. Error bars mark the standard error of the mean.*

In CA3, we found that all input types promoted R-activation of cells during sharp waves, including inhibition. The latter sounds counter-intuitive, however sharp waves are characterized by high non-rhythmic firing of both pyramidal cells and interneurons, and as such it is possible that enhanced incoming inhibitory synapses can promote post-inhibitory rebound spiking in CA3 cells rather than suppress them (Diba et al. 2014). In CA1, the results were more immediately intuitive: higher R-activation scores were found in cells with lower inhibitory and higher excitatory synaptic inputs, while a cell’s intrinsic excitability has limited influence on its R-activation (this is consistent with our analysis of CA1 cells activation across ripples in our previously developed model of CA1 ripples). This differential role of inputs in R-activation in CA3 vs CA1 is likely a result of the fundamentally different dynamics of sharp waves in CA3 and ripples in CA1, where high-frequency inhibitory rhythms leave only short windows of opportunity for pyramidal cell spiking.

**R-activation of cell pairs increases with shared excitatory input**

Beyond the activation of single cells, memories in the hippocampus are represented by ordered sequences of multiple cell spikes. The simplest way to shape spiking sequences within CA1 would be to use excitatory connections within the network, but pyramidal to pyramidal cell synapses in CA1 are few and sparse (Deuchars and Thomson 1996). Hence, we consider the larger CA3-CA1 network, within which a straightforward way to create sequences in CA1 would be to have sequences in CA3 (shaped by the highly recurrent CA3 pyramidal cells) and use the Schaffer collaterals to evoke corresponding sequences in CA1. Hence, we expect learning to shape both the CA3 recurrent connections and the CA3- to-CA1 projections. We use our model to investigate the relative role of these two main components in shaping the degree of reactivation of cell sequences.

We consider the order of spiking in cell pairs as the building block of a full sequence replay across populations (Figure 4). As for single cells, we quantified the R-activation of a cell pair as the percent of SWR in which the two cells spiked in the correct order, grouped all cell pairs with similar R-activation scores, and compared how inputs to cells in the pair changed with R-activation. We studied pairs of cells within the same region and CA3-CA1 cell pairs (Figure 4). To find meaningful inputs, capable of promoting co-spiking during a SWR, we looked for di-synaptic paths (cell A to cell B to cell C) which started from any one given cell and ended on both cells of the pair (formulae on Figure 4, details in Appendix).

**Figure 4.**
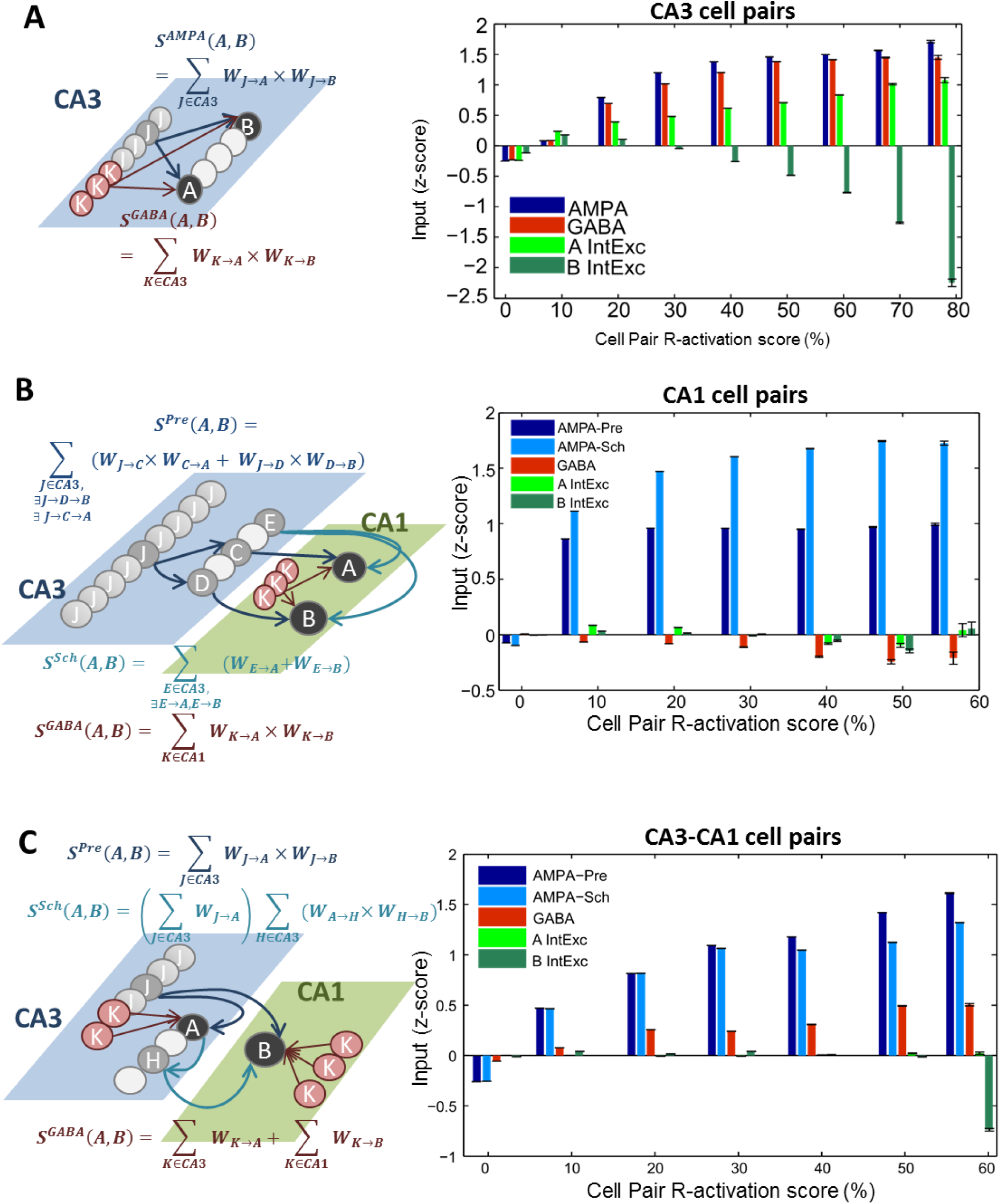
Cells that co-reactivate share their input. ***A.*** *Relationship between R-activation scores and input for pairs of CA3 pyramidal cells. Left plot: drawing and formulae introduce the synaptic AMPA and GABA inputs considered (S*^*AMPA*^ *and S*^*GABA*^*) for any cell pair labeled (A, B). These estimates quantify the amount of excitatory and inhibitory input that cells A and B have in common. Right plot: R-activation of CA3-CA3 cell pairs is related to strength of synaptic inputs and intrinsic excitability. The bar plot shows on the x –coordinate the average R-activation score of CA3-CA3 cell pairs belonging to the same score group (±5%) and on the y-coordinate the level of each different input (S*^*AMPA*^, *S*^*GABA*^ *or Intrinsic Excitability for cell A and for cell B) for cell pairs within that score group. Error bars mark the standard error of the mean. Before grouping cell pairs by their R-activation scores, each input was z-scored to enable comparisons of their respective trends. Therefore, a negative y-coordinate does not reflect a negative input, but an input below the average across the whole network.* ***B.*** *Relationship between cell pair R-activation scores and inputs for pairs of CA1 pyramidal cells. Left plot: drawing and formulae introduce the synaptic inputs considered. For cell pair (A, B) in CA1, excitatory AMPA input from Schaffer collateral alone is labeled S*^*Sch*^, *while the role of the excitability of pre-synaptic cells in CA3 is considered in defining the complementary excitatory synaptic input S*^*Pre*^. *Inhibitory synaptic input S*^*GABA*^ *is found analogously to the one for CA3-CA3 cell pairs (in panel* ***A****). These measures are introduced to quantify the shared synaptic inputs between cells A and B in each pair. Right plot: cell pair R-activation scores R-activation of CA1-CA1 cell pairs is related to strength of excitatory synaptic inputs, but not inhibitory synaptic inputs or intrinsic excitability. The bar plot shows on the x –coordinate the average R-activation score of CA1-CA1 cell pairs belonging to the same score group (±5%) and on the y-coordinate the level of each different input (S*^*Sch*^, *S*^*Pre*^, *S*^*GABA*^ *or Intrinsic Excitability for cell A and for cell B) for cell pairs within that score group. Error bars mark the standard error of the mean. Before grouping cell pairs by their R-activation scores, each input was z-scored to enable comparisons of their respective trends. Therefore, a negative y-coordinate does not reflect a negative input, but an input below the average across the whole network.* ***C.*** *Relationship between cell pair R-activation scores and inputs for pairs of CA3-CA1 pyramidal cells. Left plot: drawing and formulae introduce the synaptic inputs considered. Excitatory (AMPA-mediated) synaptic inputs are measured as S*^*Sch*^ *(which emphasize the role of synaptic paths from cell A in CA3 to cell B in CA1) and S*^*Pre*^ *(which emphasizes the role of cells in CA3 connecting to both cell A in CA3 and cell B in CA1). Inhibitory synaptic input is found in CA3 for cell A and in CA1 for cell B, the sum of the two constitutes S*^*GABA*^ *for the (A, B) cell pair. These measures are introduced to quantify the shared synaptic inputs between cells A and B in each pair. Right plot: cell pair R-activation scores R-activation of CA3-CA1 cell pairs is related to strength of excitatory synaptic inputs, partly to inhibitory synaptic inputs, but not intrinsic excitability. The bar plot shows on the x –coordinate the average R-activation score of CA3-CA1 cell pairs belonging to the same score group (±5%) and on the y-coordinate the level of each different input (S*^*Sch*^, *S*^*Pre*^, *S*^*GABA*^ *or Intrinsic Excitability for cell A and for cell B) for cell pairs within that score group. Error bars mark the standard error of the mean. Before grouping cell pairs by their R-activation scores, each input was z-scored to enable comparisons of their respective trends. Therefore, a negative y-coordinate does not reflect a negative input, but an input below the average across the whole network.*

Consistent with our single cell analyses, we considered the role of excitatory synaptic input, inhibitory synaptic input and intrinsic excitability of either one of the cells in the pair, and estimated the relative role of the different inputs on cell pair R-activation by multivariate linear regression (tables shown in Supporting Information). We found that excitatory inputs significantly modulated the R-activation scores of all cell pairs, in that pairs with higher scores received overall larger excitatory synaptic inputs. Also, we found that inhibitory synaptic inputs and intrinsic excitability of the cells in the pair significantly promoted R-activation of CA3-CA3 cell pairs, but not of CA1 cell pairs (again in agreement with what we found for single cells). In CA3-CA1 cell pairs, inhibitory synaptic inputs were found to partially contribute to R-activation, likely driven by their role on the activation of CA3 cells. Overall, this suggests a scenario in which R-activation of a spike sequence is a network-wide phenomenon, shaped by synaptic inputs differently within CA3 and CA1. It provides a framework to explore mechanisms through which experience-dependent tuning of synaptic inputs during behavior could induce changes in the R-activation seen during sleep.

### Synaptic Plasticity in both CA3 and CA1 can shape R-activation of cell pairs

To test how possible changes to synaptic paths leading to cells in a sequence could affect the R-activation of such sequence, we again studied the simplest possible sequence: a cell pair. We manually altered the synaptic paths contributing to the inputs of the cell pair (as defined in Figure 4) and compared the cell pair R-activation before and after this manipulation. Intuitively, this represents a rough approximation of what learning during awake could do to synapses: change a selected few while leaving all others unaltered. Testing this procedure on randomly selected samples of cell pairs (details in Appendix) showed that indeed such simple manipulation could promote the R-activation score of a specific cell pair without altering the overall average R-activation in the network. In other words, selective changes along di-synaptic pathways can turn a cell pair with average R-activation into a highly R-activating (memory encoding) cell pair. While it is intuitive that increasing cell excitatory inputs promotes spiking, less obvious is how spontaneous network dynamics avoid becoming catastrophically disrupted by such plasticity. For instance, SWR (and hence R-activation distributions) result from interplay between excitatory and inhibitory activity in the network, and the artificial strengthening of distributed paths in the network has the potential to lead spontaneous activity toward epileptic-like dynamics. Despite this, the average R-activation profile across all other pairs remains stable as highly reactivated pairs emerge.

Nevertheless, our analysis revealed that both connections within CA3 and projections CA3-to-CA1 are crucial in shaping the reactivation of CA1 pairs. And it predicts that by changing synaptic paths along both architectures reliability of reactivation across many SWR can be achieved. In principle, our statistical analysis over all possible pairs showed a very similar role of the two synaptic ranges (within CA3 and Schaffer collaterals), but with all possible CA1 pairs the ones with high R-activation scores are very few, so the need for both CA3-Pre and Schaffer input to shape a highly reactivating CA1 pair could be sufficient, but possibly not necessary. To test this idea, we again tested our sample CA1 pairs which received a new input (representing a simplified learning mechanism), but we only modified a synaptic path if it was from CA3 to CA1, leaving all synapses within CA3 unaltered. If only the Schaffer collaterals were necessary to mediate a strong increase in CA1 pair reactivation, we should see a similar effect as in Figure 5c.

**Figure 5.**
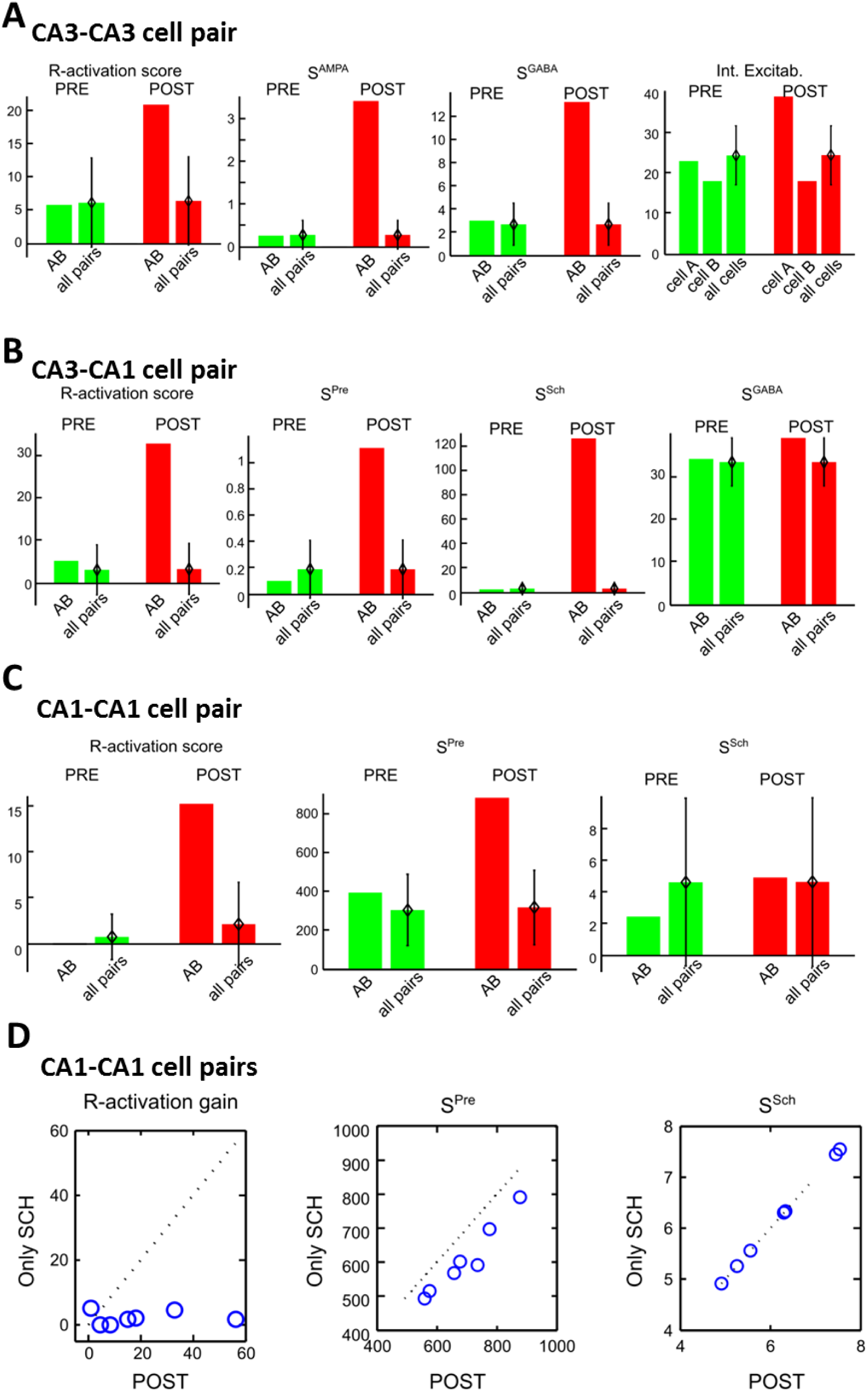
Increased synaptic strengths promote R-activation of randomly selected cell pairs. ***A.*** *One example of CA3 cell pair AB randomly chosen in a simulation. Reported on the bar plots are its R-activation score, S*^*AMPA*^, *S*^*GABA*^ *and the intrinsic excitability of both cell A and B separately. For each measured output, the value for the AB pair is shown next to a bar reporting the mean ±standard deviation across all pairs of CA3 pyramidal cells (or single cells) in the same simulation. The scaling introduced in the network connectivity and intrinsic excitability increased all considered inputs from before (PRE, green) to after (POST, red). Note that the mean and standard deviations of the inputs across the network do not change from PRE to POST. The R-activation score of AB is near the mean in PRE, and larger than one standard deviation above the mean in POST.* ***B.*** *One example of CA3-CA1 cell pair AB randomly chosen in a simulation. Reported on the bar plots are its R-activation score, S*^*Pre*^, *S*^*Sch*^ *and S*^*GABA*^. *For each measured output, the value for the AB pair is shown next to a bar reporting the mean ±standard deviation of that value across all pairs of CA3-CA1 pyramidal cells in the same simulation. The scaling introduced in the network connectivity increased all considered inputs from before (PRE, green) to after (POST, red). Note that the mean and standard deviations of the inputs across the network do not change from PRE to POST. The R-activation score of AB is near the mean in PRE, and larger than one standard deviation above the mean in POST.* ***C.*** *One example of CA1-CA1 cell pair AB randomly chosen in a simulation. Reported on the bar plots are its R-activation score, S*^*Pre*^ *and S*^*Sch*^. *For each measured output, the value for the AB pair is shown next to a bar reporting the mean ±standard deviation of that value across all pairs of CA1-CA1 pyramidal cells in the same simulation. The scaling introduced in the network connectivity increased all considered inputs from before (PRE, green) to after (POST, red). Note that the mean and standard deviations of the inputs across the network do not change from PRE to POST. The R-activation score of AB is near the mean in PRE, and larger than one standard deviation above the mean in POST.* ***D.*** *Comparing manipulating the whole CA3-CA1 architecture or only Schaffer Collaterals. For all sampled CA1-CA1 cell pairs (n=7) we compare the outcome of the general manipulation case (as in panel C, POST, x-axis in the plots) and in the case of changes introduced only in the Schaffer Collaterals (Only SCH, y-axis in the plots). If the two manipulations led to identical effects, the data points would lie along the identity line (reported in plots for reference). Note that both cases lead to the same changes in, but is smaller when no CA3-CA3 synapses are increased. The left panel shows the R-activation gain, defined as the R-activation score of the pair after the manipulation minus the one before the manipulation (in the “PRE” condition in panel C). Note that gain is much higher when the manipulation affects the complete CA3-CA3-CA1 synaptic architecture, rather than only the CA3-CA1 synapses.*

In every cell pair we tested, we found that altering only the Schaffer Collateral component of the network resulted in an identical increase of the Schaffer input and a reduced increase of the CA3-Pre input (as a reminder, the formula for this input takes into account many CA3-CA3 synapses and some CA3-to-CA1 synapses, hence the effect of the Schaffer-only manipulation on this input). While the spiking patterns in the network remained physiological under this manipulation, there were very small increases in the R-activation scores of the tested cell pairs when only the Schaffer collateral portion of the input was modified. In fact, for all pairs but one the partial manipulation led to a much smaller increase of the R-activation score for the cell pairs (see Figure 5d), compared to the increase generated by changes in both CA3 recurrents and Schaffer-collaterals. Hence, our model suggests that the CA3-CA1 synaptic architecture has to be changed as a unified system to promote reliable replay of CA1 cell pairs, which are the building blocks of full-fledged CA1 spike sequences, once again suggesting that replay in CA1 is contingent on appropriate replay in CA3.

## Discussion

In this paper, we introduced a spiking network of CA3-CA1 activity showing spontaneously emergent, localized, stochastic SWRs. Within these events, we studied the spike reactivation in CA3 and CA1 during SWRs, and measured the fraction of ripples in which a cell spiked with a “ripple-activation” (R-activation) score between 0 and 100%. When compared to a stationary sampling process, we found that a relatively rigid network architecture (defined by stationary probability distributions of intrinsic and synaptic properties) determined the spiking of low R-activating cells (the majority), while the network dynamics (dependent on the “synaptic outliers” and specific implementation of the network configuration) shaped the activity of highly R-activating cells. We further found that the degree to which a cell activated across ripples was modulated by the amount of synaptic excitatory and inhibitory input received by the cell. In particular, for CA3 cells but not for CA1 cells, we found a role for intrinsic cell excitability in shaping cell activation across ripples. This observation generalized to cell pairs and synaptic paths which impinge on both cells composing the pair, meaning that a shared pathway of synaptic input could promote co-activation of cell pairs. Increasing the shared synaptic input of a cell pair could lead to increased R-activation, and that increasing the R-activation of a pair of CA1 cells required changing synapses in both the CA3 architecture and Schaffer Collaterals. Together, these observations are indicative of a network-wide coordination of activation probability across SWRs for cells and cell pairs, which is further refined by specific synaptic strengths, such as those facilitated during awake learning. Our model predicts a continuous re-arrangement of reactivation among cell assemblies, where the balance of global network dynamics ensures the majority of cell assemblies have low reactivation, while specific synaptic strengths mediate the chance of predominant activation of a given memory during a given bout of SWR.

### Model captures both generic ripple activity and potential mechanisms of learning-dependent replay

CA1 place cells recruited during encoding of recent experience are known to reactivate together during subsequent sleep (Wilson and McNaughton 1994). Importantly, rather than displaying a uniform probability of spiking during ripples, cells can be divided in those which are active during SWRs and those which are not, a feature that persists across recordings (Buzsaki 2015; Grosmark and Buzsaki 2016). The precise manner in which *in vivo* SWRs involve or exclude a specific pyramidal cell from their activity remains unknown. Our model predicts that synaptic plasticity during learning (such as, e.g., mediated by awake SWR activity (Atherton et al. 2015; Girardeau and Zugaro 2011; O’Neill et al. 2006)) could effectively cause the inclusion of cells coding for a novel learned task in the set of CA1 pyramidal cells which are spiking during sleep SWRs (Grosmark and Buzsaki 2016). Hence, we propose that within the CA3-CA1 architecture, SWRs frame activation of a generic representation within which spikes from the specific place cells involved in recently encoded experience are preferentially engaged, gated by recent synaptic plasticity, and that this representation involves both CA3 and CA1 regions, rather than being encoded in CA1 alone.

Our analysis gives rise to a scenario in which the same network properties which enable the spontaneous emergence of SWRs in the CA3-CA1 architecture (high recurrence in CA3 pyramidal cells, noise-driven spiking in CA3, strong drive to inhibitory neurons in CA1 from CA3 activity) also select a small subset of cells which are most likely to reactivate in a high fraction of SWRs. In other words, our model predicts that replay arises by virtue of the same AMPA/GABA synaptic architecture which generates SWRs themselves. During sleep, the content of hippocampal replay can theoretically be selected within the hippocampal circuitry, and interact with cortical activity by carefully organized timing of SWRs compared to other ongoing oscillations. Within this architecture, memory formation mechanisms during wake (such as STDP, reward signals and awake replay) can modify the chances of specific cells to be replayed during sleep SWRs by altering the relative strengths of synaptic pathways impinging on a group of cells.

### Other models of SWR and hippocampal replay

The biophysical model of CA3-CA1 SWR activity which we propose in this study builds on an extensive literature on the mechanisms of ripples and sharp waves. *In vitro* and *in vivo* studies have shown that in CA1 ripples are dominated by inhibitory phasic activity (Gan et al. 2017; Schlingloff et al. 2014), and basket cells spike at high frequency (Schlingloff et al. 2014; Varga et al. 2012) in localized groups (Patel et al. 2013). Meanwhile, pyramidal cells spike relatively rarely, phase-locked to windows of opportunity left by the ongoing oscillatory inhibitory signal (Csicsvari et al. 1999a; b). In CA3, excitatory and inhibitory spiking is not locked to CA1 ripple waves (Sullivan et al. 2011), and can emerge spontaneously (Maier et al. 2003; Schlingloff et al. 2014) *in vitro*, while *in vivo* its initiation is still under investigation (Csicsvari et al. 2000; Oliva et al. 2016). In the search for explanatory mechanisms underlying SWRs, numerous possible strategies have been introduced. Gap junctions between CA3 pyramidal cells have been proposed to be necessary (Traub and Bibbig 2000; Traub et al. 1991) but experimental evidence is still not definitive for such mechanism (Buhl et al. 2003; Buzsaki 2015; Stark et al. 2014). Models which take advantage of supra-linear summation of post-synaptic potentials among CA3 and CA1 pyramidal cells have proposed that SWRs are synaptically propagating waves, where each excitatory spike induces its own local feedback inhibitory activity (Jahnke et al. 2015; Memmesheimer 2010). These assume a very similar activity in CA3 and CA1 during SWRs, and depend strictly on the presence of strong excitatory synapses between CA1 cells (which have been found to be very few (Deuchars and Thomson 1996)). In work by Taxidis et al. (Taxidis et al. 2012), AMPA and GABA receptor-mediated synaptic activity, combined with intrinsic bursting of CA3 pyramidal cells, are the basic mechanisms underlying the emergence of SWRs in a computational model which can be seen as a precursor to our model. In the model by Taxidis, SWRs have to happen rhythmically, because their initiation is crucially tied to the bursting activity in the CA3 recurrent network: that model requires that CA3 pyramidal cells spike in bursts, and do so in strong synchrony in every theta cycle. In our model, SWRs occur stochastically, with long stochastic pauses in between packets of events, consistent with *in vivo* findings (Davidson et al. 2009). This physiologically realistic result arises from taking into consideration the crucial role played by background noisy activity in setting the SWR mechanism.

In previous work (Malerba et al. 2016), we introduced a model of CA1 receiving direct current (a simplified sharp wave). In that model, ripples in CA1 were represented by transient orbits of a dynamical system in which ripple activity is initiated by a synchronizing input to interneurons, then activity winds around a fixed point inducing fast decaying oscillations, and termination is due to heterogeneity among the interneurons driving the transient orbit back to the stationary (de-synchronized) state. Here, we introduce sharp wave activity in CA3 which is an escape process. The CA3 network has strong recurrence of excitatory synapses, and CA3 pyramidal cells are in a noise-driven spiking regime, which means that spikes are driven by fluctuations in the incoming currents (including synaptic ones). This imposes a disorganized state in the network (LIA, found during slow wave sleep in the hippocampus (Buzsaki 2015)), and SWRs emerging when enough CA3 pyramidal cells spike in a small window of time. This leads part of the network to organize, accumulating recruitment of other pyramidal cells and interneuron spikes until the network cannot sustain its propagation any further. This implies that sparseness in the CA3 recursive synapse architecture is also a necessary property of our model design.

Our model design for sharp wave activity is similar to the one introduced by Omura et al. (Omura et al. 2015), which particularly addressed the lognormal distribution of firing rates found across CA3 activity and its relationship to a specific distribution in the synaptic weights of excitatory connections in the network. In their model, hippocampal activity is isolated from external input, apart from a short-lived initial Poisson drive. Our new model is a complete CA3-CA1 spontaneous activity design, where sharp waves and ripples are built to be different phenomena, one mainly excitatory, marked by wave propagation and extending to a large portion of the excitatory population, one mainly inhibitory, rhythmic and involving a small fraction of local pyramidal cells. In our model, cells receive colored noise to represent the ongoing activity of all other inputs (for example from entorhinal cortex) present *in vivo* (Atallah and Scanziani 2009). Furthermore, we focus on how this structure is capable of supporting replay mediated by AMPA and GABA synapses, which is not addressed in Omura et al. The ability of selective connections to promote cell assembly reactivation (spontaneous and evoked) has been analyzed recently by (Chenkov et al. 2017) who show that synaptic strengths among cells in one assembly can promote burst-reactivation, considering both excitatory and inhibitory cells as part of the assembly. In our study, we consider spikes of pyramidal cells to represent information content and spikes of inhibitory cells to contribute to the shape of overall network dynamics (ending a sharp wave in CA3, and pacing the frequency of ripples in CA1). This idea is consistent with experimental data which has found a heightened specificity in the activation of hippocampal pyramidal cells compared to hippocampal interneurons across the various rhythmic activities which mark different phases in information processing in an *in vivo* task (Rangel et al. 2016).

### Summary and predictions for reactivation

We believe our model is the first which addresses the mechanisms of localized activity not only in CA1 (ripples), but also in CA3 (sharp waves). Our study further expands the possibilities on how hippocampal reactivation during sleep can interact with ongoing activity in cortex and other brain structures. It predicts that topologically organized input (from CA2 or directly from mossy fibers) could selectively activate a given portion of CA3 and foster reactivation which is specific to that area (a local SWR event). The spiking content which is then reactivated (the precise spike sequence) in CA3-CA1 will depend on the specifics of synaptic connections within CA3 and between CA3 and CA1. Such replay could then be passed downstream (through subiculum and its targets) back to cortex and other structures, in an ongoing loop aimed at changing synapses outside the hippocampus based on the content of hippocampal replay activated through selective projections from upper layers of entorhinal cortex to dentate gyrus (and hence CA3). For this overall consolidation to take place, and hence perform a share-and-transfer of information from hippocampus to cortex during slow wave sleep, ripples need to be flexible in their timing, while their content needs to be stable, but able to be evoked differentially depending on the overall input activity (replay is known to change due to auditory stimulation during sleep (Carr et al. 2012), for example). Furthermore, for consolidation to take place, SWRs need to be able to reactivate recent and past events to foster the integration of new factual events in generalized conceptual schemas which enable the animal (and humans) to use its experiences to comprehend the world surrounding it (hence, generalize). A CA3-CA1 network which is too rigid, too rhythmic, or too dependent on few supra-linear connections in its specific SWR activity, will be less able to support flexible spiking to mediate consolidation across a night of sleep.

## Materials and Methods

### Network Model: rationale

We started with our previously developed (Malerba et al. 2016) network of CA1 pyramidal and basket cells and constructed a network of pyramidal and basket cells to represent CA3 activity, then built Schaffer Collaterals projecting CA3 pyramidal cells to CA1 pyramidal cells and interneurons. We used equations of the adaptive exponential integrate and fire formalism (Brette and Gerstner 2005; Touboul and Brette 2008), which can show bursting activity (like CA3 and CA1 pyramidal cells (Andersen et al. 2006)) or fast-spiking activity (like basket cells (Andersen et al. 2006)) depending on their parameters (Brette and Gerstner 2005). CA3 pyramidal cells were allowed a stronger tendency to burst in response to a current step input by having a less strong spike frequency adaptation than CA1 neurons (Andersen et al. 2006). For simplicity, all cells belonging to the same population had the same parameters (specified in the following section). To introduce heterogeneity among cells in the network, every cell received a different direct current term (selected from a normal distribution)), and every cell received an independent Ornstein–Uhlenbeck process (OU process) (Uhlenbeck and Ornstein 1930), which can be thought of as a single-pole filtered white noise, with cutoff at 100Hz. This noisy input was added to take into account the background activity of the cells which we did not explicitly model in the network. The standard deviation of the OU process controlled the size of the standard deviation in sub-threshold fluctuations of cell voltages, and was a parameter kept fixed within any cell type. Once the parameter tuning was in effect, the cells (even when disconnected from the network) were showing fast and noisy sub-threshold voltage activity, and their spikes were non-rhythmic, driven by fluctuations in the noise input they received, which is called a noise-driven spiking regime, rather than a deterministic spiking regime, and is representative of *in vivo* conditions (Broicher et al. 2012; Fernandez et al. 2011; Roxin et al. 2011).

Cells were arranged within a one-dimensional network in CA3 (see Figure 1A), and connectivity within CA3 was characterized by each cell reaching other cells within a third of the network around them (Figure 1B), which is consistent with anatomical estimates (Li et al. 1994). For pyramidal to pyramidal cells connections, the probability of synaptic contact within this radius of one third was higher for neurons closer to the pre-synaptic cell and decayed for neurons further away. Details of all network connections are introduced in the Network Model: connectivity section. Intuitively, the highly recurrent connections between pyramidal cells in CA3 had a gradient in density that resulted in a convergence/divergence connectivity fairly uniform across all CA3 pyramidal cells, which represents the overall homogeneity of CA3 pyramidal cells arborization within the region Overall, this connectivity represents the highly recurrent pyramidal connections in CA3 without introducing special hubs of increased excitatory recurrence in any specific location in the network.

### Network Model: equations and parameters

We model SWR activity in the hippocampus using a network of 240 basket cells and 1200 pyramidal cells in CA3, 160 basket cells and 800 pyramidal cells in CA1. The ratio of excitatory to inhibitory neurons is known to be approximately 4 (Andersen et al. 2006) and since in our model we did not introduce any of the numerous hippocampal interneuron types but for basket cells, we apply that ratio to the pyramidal to basket cell network. This ratio also favored the ability of the network to support a background disorganized spiking regime, where excitatory and inhibitory currents were able to balance each other (Atallah and Scanziani 2009). For each neuron, the equations are

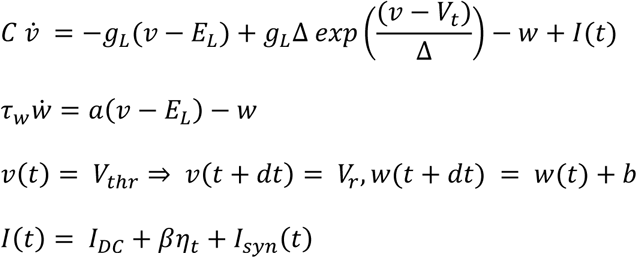

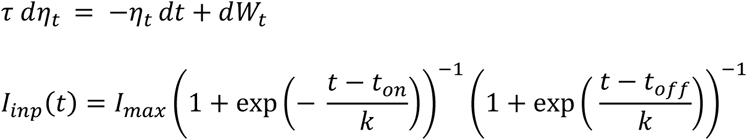

CA1 cells parameters are reported in (Malerba et al. 2016), and CA3 cells parameters were as follows. Pyramidal cells parameters: C (pF) = 200; g_L_ (nS) = 10; E_L_ (mV)= -58; A = 2; b (pA) = 40; Δ (mV) = 2; τ_w_ (ms) = 120; V_t_ (mV) = -50; V_r_ (mV) = -46; V_thr_ (mV) = 0. Interneurons parameters: C (pF) = 200; g_L_ (nS) = 10; E_L_ (mV)= -70; A = 2; b (pA) = 10; Δ (mV) = 2; τ_w_ (ms) = 30; V_t_ (mV) = -50; V_r_ (mV) = -58; V_thr_ (mV) = 0.

The coefficients establishing noise size were β = 80 for pyramidal cells, β = 90 for interneurons. DC inputs were selected from Gaussian distributions with mean 24 (pA) and standard deviation 30% of the mean for pyramidal cells in CA3, mean 130 (pA) and standard deviation 30% of the mean for CA3 interneurons, mean 40 (pA) and standard deviation 10% of the mean for CA1 pyramidal cells and mean 180 (pA) and standard deviation 10% of the mean for CA1 interneurons.

Synaptic currents were modeled with double exponential functions, for every cell *n* we had *Isyn* (*t*)=

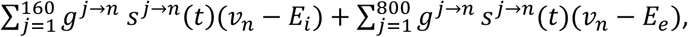

where *E*_*i*_ = –80 mV and *E*_*e*_= 0 mV, and 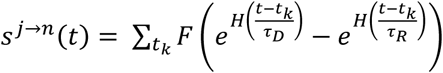, where *t*_*k*_ are all the spikes of pre-synaptic cell *j*.

In this equation, *F* is a normalization coefficient, set so that every spike in the double exponential within parentheses peaks at one, and *H*(·) is the Heaviside function, ensuring that the effect of each pre-synaptic spike affects the post-synaptic current only after the spike has happened. The time scales of rise and decay (in ms) used in the model were as follows (Bartos et al. 2002; Cutsuridis et al. 2010; Malerba et al. 2016; Taxidis et al. 2012). For AMPA connections from pyramidal cells to pyramidal cells: τ_R=_ 0.5, τ_D=_ 3.5. For AMPA connections from pyramidal cells to interneurons: τ_R=_ 0.5, τ_D=_ 3. For GABA_A_ connections from interneurons to interneurons: τ_R=_ 0.3, τ_D=_ 2. For GABA_A_ connections from interneurons to pyramidal cells: τ_R=_ 0.3, τ_D=_ 3.5.

### Network Model: connectivity

The CA3 network was organized as a one-dimensional network. For connections from a CA3 pyramidal cell to the other CA3 pyramidal cells, we first considered a radius (of about one third of the network) around the presynaptic cell, and the probability of connection from the presynaptic cell to any cell within such radius was higher for cells with indices nearby the presynaptic cell and reduced progressively with cell index distance (Li et al. 1994). Specifically, we used a cosine function to shape the probability within the radius, and parameterized how fast with index distance the probability had to decay by using a monotonic scaling of the cosine phase: if x was the index distance within the network, y = arctan(kx)/arctan(k) imposed the decay probability p(y) = Pcos(4y), where P was the peak probability and k= 2 was a parameter controlling the decay of connection probability with distance within the radius. An analogous structure underlid the probability of CA3 pyramidal cells to connect to inhibitory interneuron in CA3 and for Schaffer Collaterals to connect a CA3 pyramidal cell to CA1 pyramidal cells (Li et al. 1994). To balance the relationship between feed-forward excitation from pyramidal cells to interneurons and feedback inhibition from interneurons to pyramidal cells, probability of connection from a presynaptic basket cell to a cell within a radius (about 1/3 of the network size) was constant at 0.7, for GABA_A_ connections to both CA3 pyramidal cells and interneurons. Within CA1 connectivity was all-to-all, with the caveat that synaptic weights which were sampled at or below zero caused a removal of a given synapse. As a result, most synapses between CA1 pyramidal cells were absent, consistently with experimental findings (Deuchars and Thomson 1996). To introduce heterogeneity among synaptic connections, synaptic weights for all synapse types were sampled from Gaussian distributions with variance (σ) given by a percent of the mean (μ). Parameters used in the simulations were (we use the notation Py3 and Py1 to denote pyramidal cells in CA3 and CA1, respectively and analogously Int3/Int1 for interneurons). Py3->Py3: μ = 34, σ = 40%μ; Int3->Int3: μ = 54, σ = 40%μ; Py3->Int3: μ = 77, σ = 40%μ; Int3->Py3: μ = 55, σ = 40%μ; Py3->Py1: μ = 34, σ = 10%μ; Py3->Int1: μ = 320, σ = 10%μ; Int1->Int1: μ = 3.75, σ = 1%μ; Py1->Int1: μ = 6.7, σ = 1%μ; Int1->Py1: μ = 8.3, σ = 1%μ; Py1->Py1: μ = 0.67, σ = 1%μ. It is to note that the mean (µ) declared was normalized by the total number of cells before the variance to the mean was introduced in the distribution. Since the CA3 and CA1 networks are of different sizes, a direct comparison of the parameter values or their magnitude across regions would not account for the effective values used in the simulations.

## Appendix

### Modeling R-activation of cells with a sampling process (Figure 2)

In our model, we first generated R-activation scores across 40 simulations, each 100s long (Figure 2B). The distribution of R-activation scores in both regions showed a large positive tale, and in CA1 a fast decay. This is consistent with data suggesting that firing rates during ripples are log-normally distributed (Anastassiou et al. 2010; Buzsaki 2015; Mizuseki and Buzsaki 2013; Mizuseki et al. 2012). One model of CA3 emergent sharp wave activity suggests that this could be related to the distribution of synaptic weights used to populate the network connectivity matrices (Omura et al. 2015).

The distribution of R-activation scores tells how many cells are likely to activate in a given fraction of all SWR, while the probability of spiking (Figure 2C,D) tells if a given cell is likely to activate in many or few SWRs (e.g., p=1 would mean that a cell spike in every SWR). For each model simulation, we reported the probability of spiking in a SWR for all cells in CA3 and CA1 in Figure 2 C-D. Since in every simulation new connectivity and heterogeneity profiles were generated (see Materials and Methods, Network Model sections), we sorted the cells according to their spiking probability within SWR, from lowest to highest. Then, it was possible to find an average distribution of such probabilities, and compare between CA3 and CA1 networks. In both cases, the variance around the mean increased for cells with higher SWR activation probabilities. The larger variations for high-probability cells (right side of the plots in Figure 2 C,D) suggested that all the cells fell in one of two categories: those which fired in very few ripples (the majority) and those which fired in a large fraction of ripples (above 0.6 probability).

Across different simulations, the rules which shaped network connectivity were fixed, but the actual specific network connections changed as they were generated probabilistically. This led to measurable variations in the properties of SWR activity, such as the total count of SWR within simulation time, and the size of ripples and sharp waves (as fraction of CA3 and CA1 pyramidal cells spiking during the event). Model properties which could affect R-activation scores can be separated in stationary and dynamic. The first are the ones that depend on the average model characteristics and do not change much across simulations (such properties included the distribution of ripple frequencies, of inter-event times, and the range of reactivation scores found across the population of pyramidal cells). The second are dependent on the specific model instantiation in a given simulation (such properties included a specific cell’s spiking probability, the specific synaptic weight of each given synapse and the specific intrinsic excitability level of a given cell). To distinguish between the contributions of the stationary and network-activity dependent properties in shaping R-activation scores across simulations, we compared the distribution of R-activations that was found across simulations with a theoretical sampling model. This model was based on the average probability of spiking in CA3 and CA1 pyramidal cells (black lines in Figure 2 C-D). We sampled cells according to those probabilities, and we repeated the sampling 10,000 times and considered each sampling event a ripple in which the sampled cells had spiked. This sampling approach assumed that spiking in each SWR was memoryless and that the probability for a given cell to be spiking in any ripple was stationary (i.e. not changing in time within a simulation), so it was an extreme simplification of a complete computational model simulation. In this sampling process, the spiking of each cell across many SWR was by definition independent of spiking history and activation of other cells, and only dependent on the average probability found in simulations.

### Measuring inputs to single cells to study their role on R-activation (Figure 3)

**For CA3 pyramidal cells**, we defined the excitatory and inhibitory synaptic input (S^AMPA^ and S^GABA^, respectively, represented in Figure 3A) by summing the synaptic weights of all incoming connections to each cell. The intrinsic excitability for each CA3 pyramidal cell corresponded to the value of model parameter *I*_*DC*_, which was assigned (and different) to every cell to introduce heterogeneity among their resting potentials (see Materials and Methods, Network model: rationale). For each input, we next analyzed the distribution of the input strength across R-activation scores (Figure 3B). All inputs considered (synaptic excitatory, inhibitory and intrinsic excitability) were on average higher for cells with higher R-activation scores, implying that all the inputs could contribute to enhancing the R-activation probability of a CA3 pyramidal cell.

To study role of the potential interactions among inputs in influencing cell R-activation, statistical inference analysis was performed using multivariate linear regression (see Table S1). For each possible combination of the inputs that were considered (parameters of the model for the statistical linear fit), we found multiple fits to R-activation of CA3 pyramidal cells. When only one input at a time was considered, we found linear models of linear terms and linear models of quadratic terms. When multiple inputs were considered, we found linear models for linear terms, interactions and quadratic terms. We report in Table S1 the models, their significance level and the portion of the data they each account for. The latter is reported by means of the adjusted R^2^ terms for each model, which can be interpreted as a measure of how much of the variance in the data is captured by a given model. As can be seen from the adjusted R^2^ values in Table S1, there was a high noise component in the relationship between inputs and CA3 cells R-activation. However, most models listed showed significant contributions of the inputs considered (by p-value<0.05 criterion). For CA3, our analysis showed significant modulation of R-activation by the three inputs (synaptic excitatory, synaptic inhibitory and intrinsic excitability). It also emphasized that the influence of synaptic excitatory input was captured in the interaction with the synaptic inhibitory input and with intrinsic excitability. When considered separately, the different inputs had a similar impact on R-activation, and combining the three inputs increased their ability to represent the R-activation data.

**For CA1 pyramidal cells**, we isolated the role of the Schaffer Collateral projections by considering only the total strength of direct synaptic connections coming to a CA1 pyramidal cell from CA3 pyramidal cells (S^Sch^). Separately we analyzed the role of activation of the pre-synaptic CA3 pyramidal cells in driving post-synaptic CA1 pyramidal cells by assigning to each CA1 pyramidal cell a value S^Pre^, which characterized the average strength of the convergent CA3 inputs to all the CA3 cells that were projecting to a given CA1 cell. Synaptic inhibitory inputs to CA1 pyramidal cells were represented by S^GABA^, computed by summing the synaptic weights of all GABA connections from CA1 inhibitory neurons to each CA1 pyramidal cell.

Again, we analyzed distribution of the input strength across R-activation scores (Figure 3D). There was a clear trend towards increasing S^Pre^ and S^Sch^ for increasing R-activation of CA1 pyramidal cells, while S^GABA^ and intrinsic excitability did not show any preferential trend for increasing R-activation groups. Only for scores above 75% (very rare in the network, as can be seen in Figure 2F) it appeared that inhibitory synaptic input below the mean could favor high reactivation.

We then used fitting of a multivariate linear model. However, we first separated CA1 pyramidal cells in those with R-activation above and below a threshold of 55% (to satisfy the statistical requirements to perform linear model fitting). The model tables were populated with the same criterion used for CA3 pyramidal cells: we tried all possible input combinations with linear, quadratic and interaction terms (the latter only for statistical models with more than one input). As can be seen in Figure 2F, a small portion of all available CA1 pyramidal cells did show R-activation scores above 55%, so it is possible that this sub-group is less representative of the general influence of inputs on R-activation scores, compared to the set of cells with R-activation scores up to 55%. In studying the large cell pool (below 55%, Table S2) we found a contribution of all inputs (pre-synaptic excitatory input, Schaffer input, inhibitory input and intrinsic excitability), with a tendency for Schaffer input to contribute in relationship to the pre-synaptic excitatory input, rather than independently. When allowing for quadratic terms, the role of direct Schaffer input was rendered null, in favor of heightened influence of pre-synaptic excitatory input and synaptic inhibitory input. When the inputs were considered separately, both excitatory synaptic inputs showed the highest impact on R-activation.

In summary, the interaction of excitatory and inhibitory inputs was significant in modulating CA1 pyramidal cells R-activation, and intrinsic excitability only increased the overall impact of the multivariate model of a minor amount. Mechanistically, this implies that if synaptic activity could change only one of these inputs at a given time, it would have the largest effects on R-activation by modulating either pre-synaptic AMPA connections in CA3, or Schaffer collaterals. We point out that the limited role of intrinsic excitability on R-activation of CA1 pyramidal cells is consistent with our earlier findings on a model of CA1 ripple activity driven by current steps (Malerba et al. 2016).

### Measuring inputs to cell pairs to study their role on R-activation (Figure 4)

To extend the concept of R-activation score to cell pairs, we considered two cells and their relative order, and found in which percent of the total ripples a given cell pair spiked. Since in our definition of cell pair the order of cell spiking was considered, the R-activation scores of cell pair AB and cell pair BA were in general different. We used our R-activation score measure to group cell pairs with similar reactivation scores in the network, and studied their common synaptic inputs. Below, we introduce the details of the analysis of activation of the cell pairs and their dependency on the network inputs. We separately tested CA3-CA3, CA1-CA1 and CA3-CA1 pairs (Figure 4).

**CA3-CA3 pairs:** For pairs of two CA3 pyramidal cells (Figure 4A), we defined an excitatory synaptic input quantifier S^AMPA^, by finding all pyramidal cells in CA3 which sent synapses to both CA3 cells in the pair and considering the product of the synaptic weights from each of such cells to the cells in the pair (Figure 4A, left panel, shows a drawing of S^AMPA^). Analogously, the inhibitory input reaching a pair of CA3 pyramidal cells was quantified by S^GABA^, summing across all interneurons projecting to both pyramidal cells in the pair the product of the identified synaptic weights (again represented in Figure 4A, left panel). The intrinsic excitabilities of the two cells in the network were considered separately.

In Figure 4A, right plot, we show that both excitatory and inhibitory inputs were larger for cell pairs with higher R-activation scores. The intrinsic excitability of the first cell in the network (labeled A in the figure) also increased with R-activation of the pair, while the intrinsic excitability of the second cell in the pair (labeled B in the figure) decreased for higher R-activations. Intuitively, these opposite trends can be understood if one considers that our pair R-activation score is a measure which takes into account the order in which the two cells spiked: intrinsic excitability promotes activation of a cell but in no connection with activity of other cells. Since for a cell pair to repeat in order across ripples it is important that the second cell does not spike de-coupled from the synaptic paths which connect its activity to the first cell, having high intrinsic excitability in the second cell would hinder the R-activation of the pair.

Next, statistical inference analysis of the role played by the different inputs in establishing the R-activation of cell pairs within CA3 was performed with linear regressions (Table S3 in Supporting Information). We populated the statistical table of model analogously to the procedure for compiling the table models for single cells and inputs (in the previous Appendix section). Specifically, for each possible combination of the inputs that were considered (parameters of the model for the statistical linear fit), we found multiple fits to R-activation of CA3-CA3 pyramidal cell pairs. When only one input at a time was considered, we found linear models of linear terms and linear models of quadratic terms. When multiple inputs were considered, we found linear models for linear terms, interactions and quadratic terms. The adjusted R^2^ terms can be interpreted as a measure of how much of the variance in the data is captured by a given model. As can be seen from the adjusted R^2^ values, there was a high noise component in the relationship between inputs and cell pair R-activation. However, the models listed showed significant contributions of the inputs considered (by p-value<0.05 criterion). The complete list of models found is reported in Table S3.

This analysis revealed a significant contribution of all inputs and their interactions to the R-activation of cell pairs within CA3. When considering subgroups of inputs, excitatory synaptic input and the intrinsic excitability of the first cell in the pair (and their interaction) could account for a large portion of the modulation of R-activation by all inputs (and interactions). When taken separately, synaptic inputs (both excitatory and inhibitory) still retained an impact on shaping the R-activation of cell pairs, however intrinsic excitability of individual cells had a very small impact on R-activation of CA3-CA3 cell pairs.

**CA1-CA1 pairs:** We extended the analysis to pairs of CA1 pyramidal cells, by introducing quantifiers of excitatory and inhibitory inputs. Inhibitory input S^GABA^ was calculated in the same manner as for pairs of CA3 pyramidal cells, meaning we found CA1 inhibitory interneurons projecting to both CA1 pyramidal cells in the pair, multiplied the synaptic strength and summed across all interneurons impinging on both cells in the pair (a representation is shown in Figure 4B, left panel). To quantify the excitatory synaptic inputs reaching a pair of CA1 pyramidal cells, we defined two separate inputs: one considering only the Schaffer Collateral contribution (S^Sch^) and one emphasizing the role of synaptic paths within CA3 ultimately reaching the cell pair in CA1 (S^Pre^). The inputs driven by the sole Schaffer Collaterals (S^Sch^) were quantified by finding cells in CA3 which projected to both cells in the pair (in CA1). The synaptic weights reaching the two cells in the pair were then summed, and these quantities were further summed across all the CA3 pre-synaptic cells found (formula and a drawing are introduced in Figure 4B left panel). The role of CA3 connectivity on the activity of a CA1-CA1 cell pair (S^Pre^) was quantified by assigning to each cell pair a value, found as follows. We first found paths of two subsequent synapses (di-synaptic paths, from a cell to the next, to the next) starting from one cell in CA3 and terminating on both cells of the CA1 pair (cells in the middle of the di-synaptic paths had to be CA3 pyramidal cells). These initial CA3 cells could drive spiking which impinged (in two synapses) on both cells of the CA1-CA1 cell pair. The synaptic weights along the paths found this way were combined, and further summed across all the possible di-synaptic paths from CA3 to CA1 found in the network (the formula is shown on Figure 4B, left panel). The magnitudes of intrinsic excitability for the two cells in the pair were considered separately.

Statistical inference analysis (Table S4) was performed analogously to the method for CA3-CA3 cell pairs. It showed that the synaptic excitatory inputs and their interactions strongly affected R-activation scores of CA1-CA1 pyramidal cell pairs. Inhibitory synaptic input and intrinsic excitability of the cells in the pair did score significantly in the model, but increased minimally the overall ability of the model to represent R-activations. In other words, including inhibitory inputs and intrinsic excitability added great complexity to the model without making significant progress on the representation of R-activations as function of the inputs. Hence, excitatory synaptic inputs greatly dominated all other deterministic inputs in shaping CA1-CA1 cell pair R-activations.

**CA3-CA1 pairs:** Since SWR dynamics were organized in a CA3 excitatory event inducing inhibitory oscillations in CA1, we considered only ordered cell pairs in which the first cell was from CA3 and the second from CA1. To define quantifiers for synaptic inputs to pairs of CA3-CA1 pyramidal cells, we again looked for synaptic paths connecting both cells in the pair (showed in Figure 4C, left panel). One direct synaptic input (S^Pre^), considered CA3 pyramidal cells that projected synapses onto both cells in the pair, and multiplied the two synaptic weights, and summed across all found pre-synaptic cells. Another excitatory synaptic input quantifier (S^Sch^) was shaped to identify synaptic paths from the first cell in the pair (in CA3) to the second cell in the pair (in CA1), by finding all di-synaptic paths from the first cell in the pair to the second cell in the pair, multiplying the synaptic weights found along the paths and scaling the resulting quantity by the total excitatory synaptic input reaching the first cell in the pair (formula and drawing in Figure 4C). Since each pyramidal cell in the pair could receive inhibition only from interneurons within its same region, the inhibitory synaptic inputs to the two cells in the pair were quantified first separately following the definitions used for single cells in CA3 and in CA1, and the pair inhibitory synaptic input S^GABA^ was computed by the sum of their respective inputs. Intrinsic excitability for each cell in the pair was considered separately.

To describe the differential role of inputs in shaping the R-activation of CA3-CA1 cell pairs, we derived a statistical inference analysis using multiple variable linear regressions (Table S5, populated with the same method as Tables S4 and S3). We found a strong dominance of excitatory synaptic inputs over the R-activation scores of CA3-CA1 pairs. When considered separately, pre-synaptic and Schaffer AMPA inputs both accounted for most of the impact of inputs on R-activation, in comparison to the effect found when considering all inputs and all their interactions. In particular, the Schaffer input had the strongest independent impact on R-activations. Furthermore, intrinsic excitability of either cell in the pair, while qualifying for significance in affecting the R-activation score, again did not introduce any strong improvement on the ability of excitatory synaptic inputs (and their interactions) to shape R-activations of CA3-CA1 cell pairs. In summary, what was true for single CA1 pyramidal cells and CA1 cell pairs carries over to CA3-CA1 cells pairs, emphasizing the dominant role of paths of excitatory synaptic connections over inhibitory ones and intrinsic excitability in promoting activation of ordered cell pairs across many SWRs.

### Altering inputs to selected cell pairs to test learning effect on subsequent R-activation (Figure 5)

We tested whether changing inputs (as defined in Figure 4) to a randomly chosen cell pair did in fact result in an increase in its R-activation score. We started by randomly choosing the cell pair, labeled A and B. In a first set of simulations (representing, for example, the sleep on the night before a learning experience, marked with PRE in Figure 5) we found the R-activation score for cell pair AB, together with the deterministic synaptic inputs to AB and intrinsic excitability of A and B. Depending on the type of cell pair considered (CA3-CA3, CA3-CA1 or CA1-CA1) we chose which deterministic inputs were likely to be most impactful on the R-activation score of the cell pair, based on our finding in Figure 4. We then re-scaled the strengths of all synaptic connections which contributed to the chosen inputs (and the parameter controlling intrinsic excitability where appropriate), so that in the new connectivity profile the cell pair AB would have larger inputs. We next ran a new simulation, to test how the spontaneous R-activation of the cell pair AB would change for increasing inputs. It is to note that we require that our manipulation preserved the main properties of the spontaneous SWR activity in the network within physiological bounds (i.e. the network did not show constantly firing cells, or highly rhythmically occurring SWRs).

Specifically, for pairs of CA3 pyramidal cells, we had previously found (Figure 4A) that excitatory and inhibitory synaptic inputs and intrinsic excitability of the first cell of the pair were larger for higher R-activating cell pairs. Hence, for a randomly chosen pair AB, we scaled synaptic connections contributing to S^AMPA^(A,B) and S^GABA^(A,B), and increased the intrinsic excitability of A. The scaling was uniform across all synapses contributing to the inputs, and it was cell pair specific, because its specific value was derived by requiring that the inputs considered will increase at least one standard deviation above the network mean after scaling. As a result, a small percentage of AMPA and GABA synapses within the CA3 network was scaled (less than 0.6% of AMPA and less than 2% of GABA synapses). This led to an increase of the cell pair R-activation score from mean value of approximately 5% to about 20% (more than one standard deviation above the mean, shown in the leftmost bar plot of Figure 5A). In a total of 6 tests of randomly selected cell pairs and simulations, the change of selected inputs and excitability led to increased cell pair R-activation score, while maintaining a network activity profile well within physiological bounds. Hence, for CA3-CA3 pairs, we concluded that uniform scaling of all synapses co-impinging on a pair could promote R-activation of that pair.

For CA3-CA1 cell pairs, we chose to modify both excitatory (S^Pre^ and S^Sch^) and inhibitory (S^GABA^) synaptic inputs, since they all showed and increasing trend for increasing R-activation of CA3-CA1 cell pairs (Figure 4C). Hence, our scaling involved AMPA synapses within CA3 and from CA3 to CA1 pyramidal cells, and GABA synapses within CA3 and within CA1. In one example of a randomly selected CA3-CA1 cell pair AB, shown in Figure 5B, the synaptic manipulation resulted in increased excitatory and inhibitory synaptic inputs on cell pair AB, while the mean and standard deviation of each input was not altered (the change affected less than 2% of AMPA synapses within CA3, less than 0.5% of Schaffer collaterals, less than 3% of GABA synapses in CA3 and about 12% of GABA synapses in CA1). The change in synapses produced a significant increase of AB R-activation score, from ∼7% to more than 30%. Among a total of 6 randomly selected CA3-CA1 cell pairs and simulations, analogous manipulations resulted in increased R-activation score for the cell pair in 4 cases, while all tests showed SWR activity within physiological bounds.

Finally, to study the effect of synaptic scaling on the R-activation of a CA1-CA1 cell pair, we elected to modify only the excitatory synaptic inputs reaching the cell pair (S^Pre^ and S^Sch^, defined in Figure 4B), since inhibitory synaptic inputs and intrinsic excitability of either cell in the pair did not show a clear increasing trend for increasing R-activation score across CA1-CA1 cell pairs (Figure 4B). In one example of randomly selected AB CA1-CA1 cell pair (Figure 5C), both excitatory synaptic inputs increased due to our synaptic scaling procedure, and AB R-activation score grew from about 2% to above 15%. In contrast to other types of cell pairs, we found that for CA1-CA1 cell pairs the scaling of excitatory synaptic inputs very often affected the network dynamics. In a total of 11 randomly selected cell pairs and simulations which resulted in increased AB R-activation, most of them (9 samples) showed an exaggerated amount of SWRs in network activity following synaptic scaling. This was likely due to the much larger fraction of excitatory synapses being modified by the scaling procedure (about 5% of all AMPA synapses between CA3 pyramidal cells, and 0.05% of Schaffer collaterals) compared to other types of cell pairs. To avoid this pitfall, we studied the complementary problem: whether reducing the input to a randomly selected CA1-CA1 cell pair would cause a reduction in the cell pair R-activation score. In a total of 7 randomly selected cell pairs and simulations, we scaled synapses within CA3 pyramidal cells and from CA3 pyramidal cells to CA1 pyramidal cells to reduce the S^Pre^ and S^Sch^ on pair AB. As expected, in all tests the network activity remained physiological, and we found in 5 tests that the synaptic manipulation resulted in lower R-activation score for the selected cell pair.

For the last group of 7 cell pairs tested, we repeated the test but only modified a synaptic strength if it belonged to Schaffer Collaterals, i.e. if it connected a CA3 to a CA1 pyramidal cell. The change we applied was identical to the one in the complete test. This led to the data points shown in Figure 5D, where we compare “POST” values (the ones from the complete test) and “Only-SCH” values (the ones from this new test).

## Acknowledgments

PM and MB work supported by ONR grant (MURI: N000141612829 and N000141612415) to MB, MJ work supported by A Wellcome Trust Senior Research Fellowship in Basic Biomedical Science 202810/Z/16/Z. The authors thank Dr. Giri Krishnan for helpful discussions.

## Supporting Information

**Table S1.** Linear regression models for R-activation of CA3 pyramidal cells vs inputs. For each possible combination of the inputs considered (parameters of the model for the statistical linear fit), we found multiple fits to R-activation of CA3 pyramidal cells. When only one input at a time was considered, we found linear models of linear terms and linear models of quadratic terms. When multiple inputs were considered, we found linear models for linear terms, interactions and quadratic terms. The models found are listed in the middle column, the value 1 represents the intercept at the origin, x1 represents the first input listed, x2 the second and so on. The adjusted R^2^ terms can be interpreted as a measure of how much of the variance in the data is captured by a given model. As can be seen from the values, there was a high noise component in the relationship between inputs and CA3 cells R-activation. However, all models listed showed significant contributions of the inputs considered (by p-value<0.05 criterion). The highlighted lines mark models used to determine the relative relevance of different inputs on CA3 R-activation.

| Inputs Considered | Model Found | Adjusted R<sup>2</sup> |
| --- | --- | --- |
| AMPA,GABA,DC | 1+x1+x2+x3 | 0.247 |
| AMPA,GABA,DC | 1+x2+x1x3 | 0.248 |
| AMPA,GABA,DC | 1+x1x2+x1x3+x2<sup>2</sup>+x3<sup>2</sup> | 0.249 |
| AMPA,GABA | 1+x1+x2 | 0.157 |
| AMPA,GABA | 1+x1+x3 | 0.157 |
| AMPA,GABA | 1+x1x2+x2<sup>2</sup> | 0.158 |
| AMPA,DC | 1+x1+x2 | 0.237 |
| AMPA,DC | 1+x1x2 | 0.237 |
| AMPA,DC | 1+x1x2+x2<sup>2</sup> | 0.238 |
| GABA,DC | 1+x1+x2 | 0.185 |
| GABA,DC | 1+x1x2 | 0.185 |
| GABA,DC | 1+x1x2+x1<sup>2</sup>+x2<sup>2</sup> | 0.2 |
| AMPA | 1+x1 | 0.146 |
| AMPA | 1+x1 | 0.146 |
| GABA | 1+x1 | 0.095 |
| GABA | 1+x1+x1<sup>2</sup> | 0.11 |
| DC | 1+x1 | 0.0906 |
| DC | 1+x1+x1<sup>2</sup> | 0.0907 |

**Table S2.**
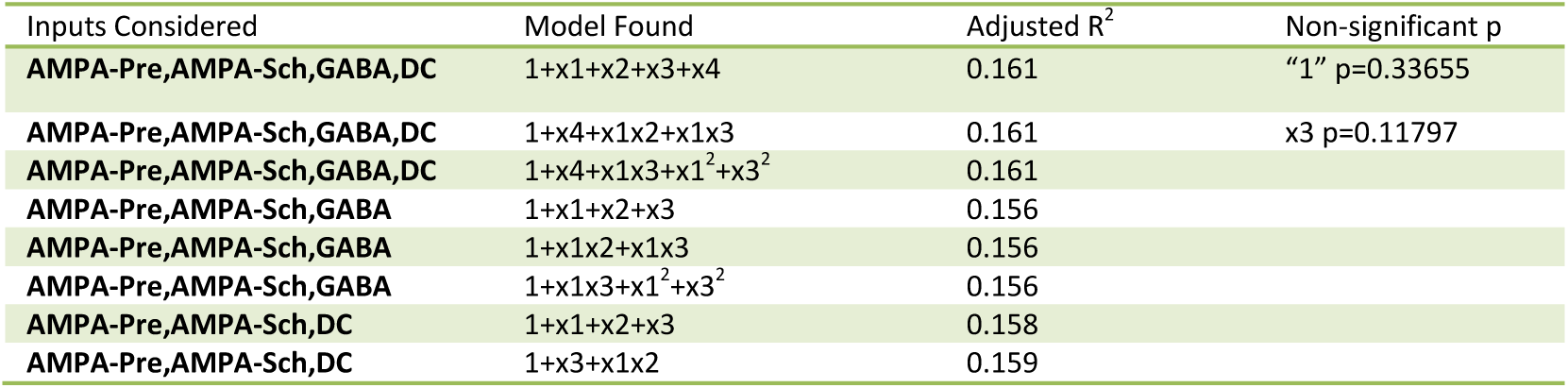
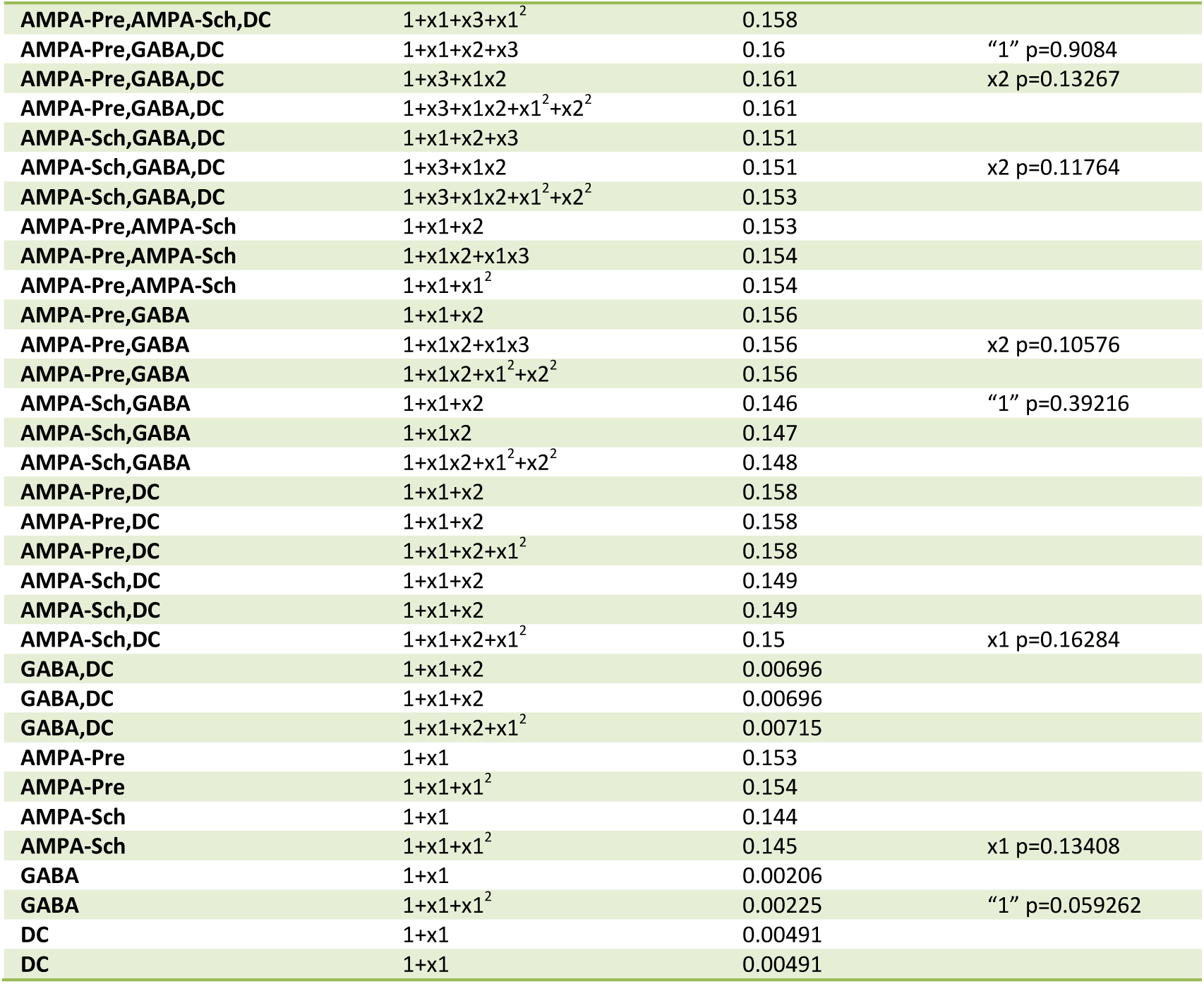
Linear regression models for R-activation of CA1 pyramidal cells(below 55% R-activation) vs inputs. For each possible combination of the inputs considered (parameters of the model for the statistical linear fit), we found multiple fits to R-activation of CA1 pyramidal cells. When only one input at a time was considered, we found linear models of linear terms and linear models of quadratic terms. When multiple inputs were considered, we found linear models for linear terms, interactions and quadratic terms. The models found are listed in the middle column, the value 1 represents the intercept at the origin, x1 represents the first input listed, x2 the second and so on. On occasion, one parameter in a given model did not pass the p-value significance requirement (p<0.05), in which case, it was declared in the last column. The adjusted R2 terms can be interpreted as a measure of how much of the variance in the data is captured by a given model. As can be seen from the values, there is a high noise component in the relationship between inputs and CA1 cells R-activation. However, the vast majority of models listed showed significant contributions of the inputs considered (by p-value<0.05 criterion).

**Table S3.** Linear regression models for R-activation of CA3-CA3 pyramidal cell pairs vs inputs. For each possible combination of the inputs considered (parameters of the model for the statistical linear fit), we found multiple fits to R-activation of CA3-CA3 pyramidal cell pairs. When only one input at a time was considered, we found linear models of linear terms and linear models of quadratic terms. When multiple inputs were considered, we found linear models for linear terms, interactions and quadratic terms. The models found are listed in the middle column, the value 1 represents the intercept at the origin, x1 represents the first input listed, x2 the second and so on. On no occasion, parameters in the models did not pass the p-value significance requirement (p<0.05; hence, the table does not need a p-value last column. The adjusted R2 terms can be interpreted as a measure of how much of the variance in the data is captured by a given model. As can be seen from the values, there is a high noise component in the relationship between inputs and cell pair R-activation. However, the vast majority of models listed showed significant contributions of the inputs considered (by p-value<0.05 criterion).

| Inputs Considered | Model Found | Adjusted $R^2$ |
| --- | --- | --- |
| AMPA-Pre,GABA,Dca,DCb | $1+x_1+x_2+x_3+x_4$ | 0.239 |
| AMPA-Pre,GABA,Dca,DCb | $1+x_1x_2+x_1x_3+x_1x_4*x_2x_3+x_2x_4+x_3x_4$ | 0.25 |
| AMPA-Pre,GABA,Dca,DCb | $1+x_1+x_2+x_3+x_4+x_1x_2+x_1x_3+x_1x_4+x_2x_3+x_2x_4+x_3x_4+x_1^2+x_2^2+x_3^2+x_4^2$ | 0.258 |
| AMPA-Pre,GABA,Dca | $1+x_1+x_2+x_3$ | 0.235 |
| AMPA-Pre,GABA,Dca | $1+x_1x_2+x_1x_3+x_2x_3$ | 0.24 |
| AMPA-Pre,GABA,Dca | $1+x_1x_2+x_1x_3+x_2x_3+x_1^2+x_2^2+x_3^2$ | 0.2479 |
| AMPA-Pre,GABA,DCb | $1+x_1+x_2+x_3$ | 0.173 |
| AMPA-Pre,GABA,DCb | $1+x_1x_2+x_1x_3+x_2x_3$ | 0.18 |
| AMPA-Pre,GABA,DCb | $1+x_1x_2+x_1x_3+x_2x_3+x_1^2+x_2^2+x_3^2$ | 0.187 |
| AMPA-Pre,Dca,DCb | $1+x_1+x_2+x_3$ | 0.225 |
| AMPA-Pre,Dca,DCb | $1+x_1x_2+x_1x_3+x_2x_3$ | 0.234 |
| AMPA-Pre,Dca,DCb | $1+x_1x_2+x_1x_3+x_2x_3+x_1^2+x_2^2+x_3^2$ | 0.244 |
| GABA,Dca,DCb | $1+x_1+x_2+x_3$ | 0.193 |
| GABA,Dca,DCb | $1+x_1x_2+x_1x_3+x_2x_3$ | 0.202 |
| GABA,Dca,DCb | $1+x_1x_2+x_1x_3+x_2x_3+x_1^2+x_2^2+x_3^2$ | 0.203 |
| AMPA-Pre,GABA | $1+x_1+x_2$ | 0.169 |
| AMPA-Pre,GABA | $1+x_1x_2$ | 0.171 |
| AMPA-Pre,GABA | $1+x_1x_2+x_1^2+x_2^2$ | 0.179 |
| AMPA-Pre,Dca | $1+x_1+x_2$ | 0.221 |
| AMPA-Pre,Dca | $1+x_1x_2$ | 0.224 |
| AMPA-Pre,Dca | $1+x_1x_2+x_1^2+x_2^2$ | 0.234 |
| GABA,Dca | $1+x_1+x_2$ | 0.19 |
| GABA,Dca | $1+x_1x_2$ | 0.193 |
| GABA,Dca | $1+x_1x_2+x_1^2+x_2^2$ | 0.194 |
| AMPA-Pre,DCb | $1+x_1+x_2$ | 0.16 |
| AMPA-Pre,DCb | $1+x_1x_2$ | 0.164 |
| AMPA-Pre,DCb | $1+x_1x_2+x_1^2+x_2^2$ | 0.174 |
| GABA,DCb | $1+x_1+x_2$ | 0.128 |
| GABA,DCb | $1+x_1x_2$ | 0.132 |
| GABA,DCb | $1+x_1x_2+x_1^2+x_2^2$ | 0.133 |
| Dca,DCb | $1+x_1+x_2$ | 0.0696 |
| Dca,DCb | $1+x_1x_2$ | 0.0705 |
| Dca,DCb | $1+x_1x_2+x_1^2+x_2^2$ | 0.0707 |
| AMPA-Pre | $1+x_1$ | 0.156 |
| AMPA-Pre | $1+x_1+x_1^2$ | 0.165 |
| GABA | $1+x_1$ | 0.124 |
| GABA | $1+x_1+x_1^2$ | 0.125 |
| Dca | $1+x_1$ | 0.0657 |
| Dca | $1+x_1+x_1^2$ | 0.0658 |
| DCb | $1+x_1$ | 0.00392 |
| DCb | $1+x_1+x_1^2$ | 0.00393 |

**Table S4.**
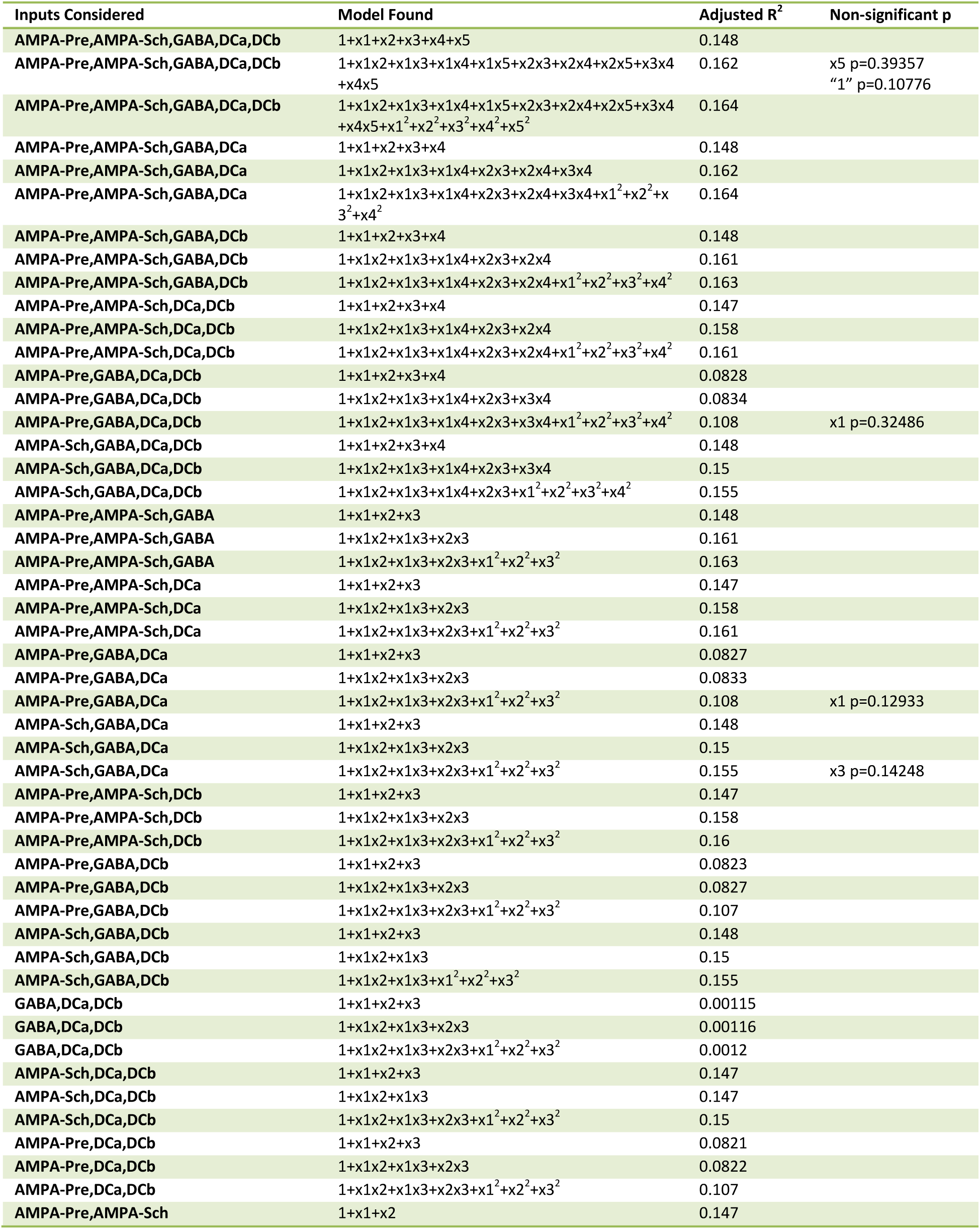
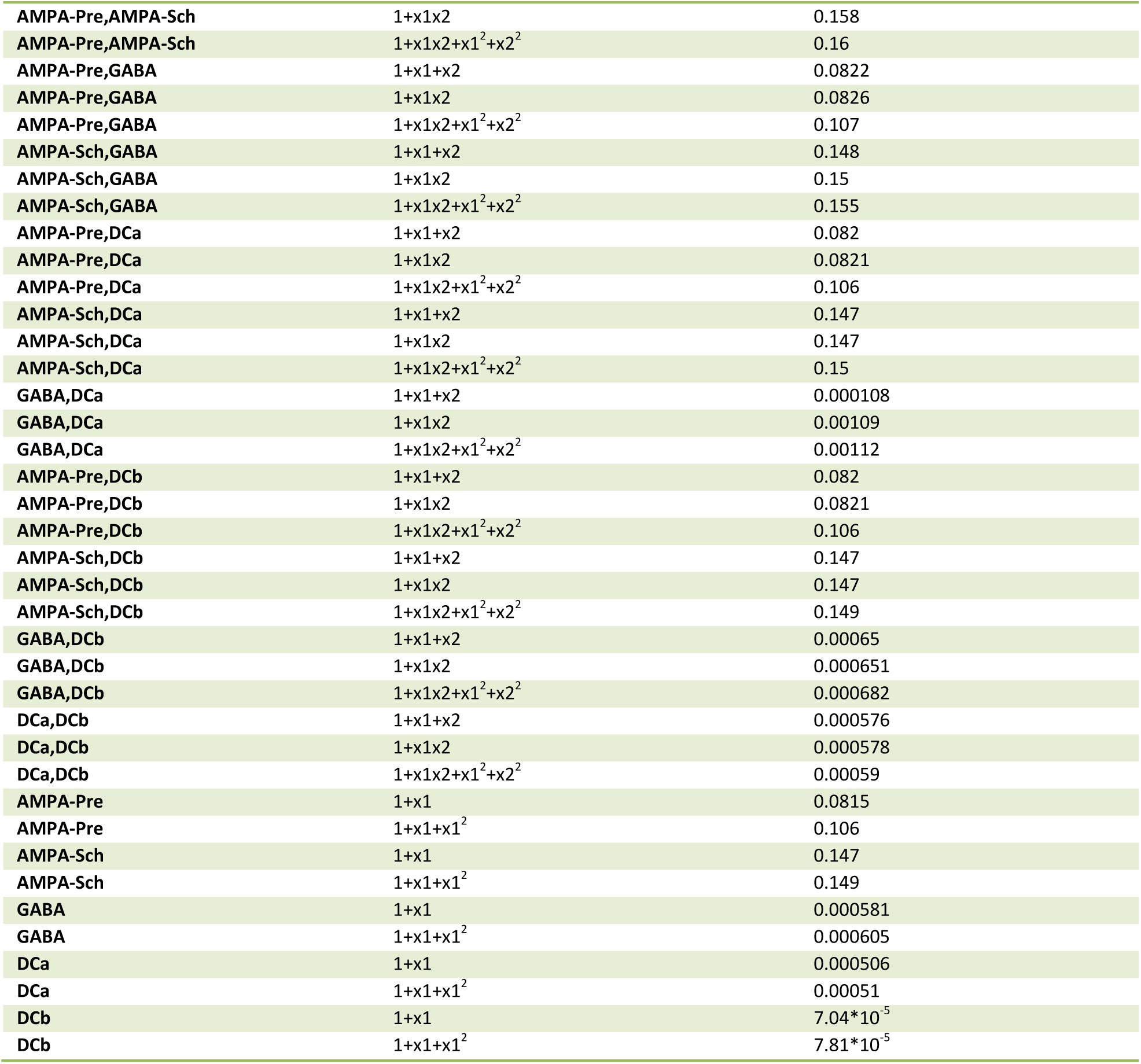
Linear regression models for R-activation of CA1-CA1 pyramidal cell pairs vs inputs. For each possible combination of the inputs considered (parameters of the model for the statistical linear fit), we found multiple fits to R-activation of CA1-CA1 pyramidal cell pairs. When only one input at a time was considered, we found linear models of linear terms and linear models of quadratic terms. When multiple inputs were considered, we found linear models for linear terms, interactions and quadratic terms. The models found are listed in the middle column, the value 1 represents the intercept at the origin, x1 represents the first input listed, x2 the second and so on. On occasion, parameters in the models did not pass the p-value significance requirement (p<0.05), in which case they were listed on the p-value last column. The adjusted R2 terms can be interpreted as a measure of how much of the variance in the data is captured by a given model. As can be seen from the values, there is a high noise component in the relationship between inputs and cell pair R-activation. However, the vast majority of models listed showed significant contributions of the inputs considered (by p-value<0.05 criterion).

| Inputs Considered | Model Found | Adjusted R<sup>2</sup> | Non-significant p |
| --- | --- | --- | --- |
| AMPA-Pre,AMPA-Sch,GABA,Dca,DCb | 1+x1+x2+x3+x4+x5 | 0.148 |  |
| AMPA-Pre,AMPA-Sch,GABA,Dca,DCb | 1+x1x2+x1x3+x1x4+x1x5+x2x3+x2x4+x2x5+x3x4+x4x5 | 0.162 | x5 p=0.39357<br>"1" p=0.10776 |
| AMPA-Pre,AMPA-Sch,GABA,Dca,DCb | 1+x1x2+x1x3+x1x4+x1x5+x2x3+x2x4+x2x5+x3x4+x4x5+x1<sup>2</sup>+x2<sup>2</sup>+x3<sup>2</sup>+x4<sup>2</sup>+x5<sup>2</sup> | 0.164 |  |
| AMPA-Pre,AMPA-Sch,GABA,Dca | 1+x1+x2+x3+x4 | 0.148 |  |
| AMPA-Pre,AMPA-Sch,GABA,Dca | 1+x1x2+x1x3+x1x4+x2x3+x2x4+x3x4 | 0.162 |  |
| AMPA-Pre,AMPA-Sch,GABA,Dca | 1+x1x2+x1x3+x1x4+x2x3+x2x4+x3x4+x1<sup>2</sup>+x2<sup>2</sup>+x3<sup>2</sup>+x4<sup>2</sup> | 0.164 |  |
| AMPA-Pre,AMPA-Sch,GABA,DCb | 1+x1+x2+x3+x4 | 0.148 |  |
| AMPA-Pre,AMPA-Sch,GABA,DCb | 1+x1x2+x1x3+x1x4+x2x3+x2x4 | 0.161 |  |
| AMPA-Pre,AMPA-Sch,GABA,DCb | 1+x1x2+x1x3+x1x4+x2x3+x2x4+x1<sup>2</sup>+x2<sup>2</sup>+x3<sup>2</sup>+x4<sup>2</sup> | 0.163 |  |
| AMPA-Pre,AMPA-Sch,Dca,DCb | 1+x1+x2+x3+x4 | 0.147 |  |
| AMPA-Pre,AMPA-Sch,Dca,DCb | 1+x1x2+x1x3+x1x4+x2x3+x2x4 | 0.158 |  |
| AMPA-Pre,AMPA-Sch,Dca,DCb | 1+x1x2+x1x3+x1x4+x2x3+x2x4+x1<sup>2</sup>+x2<sup>2</sup>+x3<sup>2</sup>+x4<sup>2</sup> | 0.161 |  |
| AMPA-Pre,GABA,Dca,DCb | 1+x1+x2+x3+x4 | 0.0828 |  |
| AMPA-Pre,GABA,Dca,DCb | 1+x1x2+x1x3+x1x4+x2x3+x3x4 | 0.0834 |  |
| AMPA-Pre,GABA,Dca,DCb | 1+x1x2+x1x3+x1x4+x2x3+x3x4+x1<sup>2</sup>+x2<sup>2</sup>+x3<sup>2</sup>+x4<sup>2</sup> | 0.108 | x1 p=0.32486 |
| AMPA-Sch,GABA,Dca,DCb | 1+x1+x2+x3+x4 | 0.148 |  |
| AMPA-Sch,GABA,Dca,DCb | 1+x1x2+x1x3+x1x4+x2x3+x3x4 | 0.15 |  |
| AMPA-Sch,GABA,Dca,DCb | 1+x1x2+x1x3+x1x4+x2x3+x1<sup>2</sup>+x2<sup>2</sup>+x3<sup>2</sup>+x4<sup>2</sup> | 0.155 |  |
| AMPA-Pre,AMPA-Sch,GABA | 1+x1+x2+x3 | 0.148 |  |
| AMPA-Pre,AMPA-Sch,GABA | 1+x1x2+x1x3+x2x3 | 0.161 |  |
| AMPA-Pre,AMPA-Sch,GABA | 1+x1x2+x1x3+x2x3+x1<sup>2</sup>+x2<sup>2</sup>+x3<sup>2</sup> | 0.163 |  |
| AMPA-Pre,AMPA-Sch,Dca | 1+x1+x2+x3 | 0.147 |  |
| AMPA-Pre,AMPA-Sch,Dca | 1+x1x2+x1x3+x2x3 | 0.158 |  |
| AMPA-Pre,AMPA-Sch,Dca | 1+x1x2+x1x3+x2x3+x1<sup>2</sup>+x2<sup>2</sup>+x3<sup>2</sup> | 0.161 |  |
| AMPA-Pre,GABA,Dca | 1+x1+x2+x3 | 0.0827 |  |
| AMPA-Pre,GABA,Dca | 1+x1x2+x1x3+x2x3 | 0.0833 |  |
| AMPA-Pre,GABA,Dca | 1+x1x2+x1x3+x2x3+x1<sup>2</sup>+x2<sup>2</sup>+x3<sup>2</sup> | 0.108 | x1 p=0.12933 |
| AMPA-Sch,GABA,Dca | 1+x1+x2+x3 | 0.148 |  |
| AMPA-Sch,GABA,Dca | 1+x1x2+x1x3+x2x3 | 0.15 |  |
| AMPA-Sch,GABA,Dca | 1+x1x2+x1x3+x2x3+x1<sup>2</sup>+x2<sup>2</sup>+x3<sup>2</sup> | 0.155 | x3 p=0.14248 |
| AMPA-Pre,AMPA-Sch,DCb | 1+x1+x2+x3 | 0.147 |  |
| AMPA-Pre,AMPA-Sch,DCb | 1+x1x2+x1x3+x2x3 | 0.158 |  |
| AMPA-Pre,AMPA-Sch,DCb | 1+x1x2+x1x3+x2x3+x1<sup>2</sup>+x2<sup>2</sup>+x3<sup>2</sup> | 0.16 |  |
| AMPA-Pre,GABA,DCb | 1+x1+x2+x3 | 0.0823 |  |
| AMPA-Pre,GABA,DCb | 1+x1x2+x1x3+x2x3 | 0.0827 |  |
| AMPA-Pre,GABA,DCb | 1+x1x2+x1x3+x2x3+x1<sup>2</sup>+x2<sup>2</sup>+x3<sup>2</sup> | 0.107 |  |
| AMPA-Sch,GABA,DCb | 1+x1+x2+x3 | 0.148 |  |
| AMPA-Sch,GABA,DCb | 1+x1x2+x1x3 | 0.15 |  |
| AMPA-Sch,GABA,DCb | 1+x1x2+x1x3+x1<sup>2</sup>+x2<sup>2</sup>+x3<sup>2</sup> | 0.155 |  |
| GABA,Dca,DCb | 1+x1+x2+x3 | 0.00115 |  |
| GABA,Dca,DCb | 1+x1x2+x1x3+x2x3 | 0.00116 |  |
| GABA,Dca,DCb | 1+x1x2+x1x3+x2x3+x1<sup>2</sup>+x2<sup>2</sup>+x3<sup>2</sup> | 0.0012 |  |
| AMPA-Sch,Dca,DCb | 1+x1+x2+x3 | 0.147 |  |
| AMPA-Sch,Dca,DCb | 1+x1x2+x1x3 | 0.147 |  |
| AMPA-Sch,Dca,DCb | 1+x1x2+x1x3+x2x3+x1<sup>2</sup>+x2<sup>2</sup>+x3<sup>2</sup> | 0.15 |  |
| AMPA-Pre,Dca,DCb | 1+x1+x2+x3 | 0.0821 |  |
| AMPA-Pre,Dca,DCb | 1+x1x2+x1x3+x2x3 | 0.0822 |  |
| AMPA-Pre,Dca,DCb | 1+x1x2+x1x3+x2x3+x1<sup>2</sup>+x2<sup>2</sup>+x3<sup>2</sup> | 0.107 |  |
| AMPA-Pre,AMPA-Sch | 1+x1+x2 | 0.147 |  |

|  |  |  |
| --- | --- | --- |
| AMPA-Pre,AMPA-Sch | $1+x_1x_2$ | 0.158 |
| AMPA-Pre,AMPA-Sch | $1+x_1x_2+x_1^2+x_2^2$ | 0.16 |
| AMPA-Pre,GABA | $1+x_1+x_2$ | 0.0822 |
| AMPA-Pre,GABA | $1+x_1x_2$ | 0.0826 |
| AMPA-Pre,GABA | $1+x_1x_2+x_1^2+x_2^2$ | 0.107 |
| AMPA-Sch,GABA | $1+x_1+x_2$ | 0.148 |
| AMPA-Sch,GABA | $1+x_1x_2$ | 0.15 |
| AMPA-Sch,GABA | $1+x_1x_2+x_1^2+x_2^2$ | 0.155 |
| AMPA-Pre,DCa | $1+x_1+x_2$ | 0.082 |
| AMPA-Pre,DCa | $1+x_1x_2$ | 0.0821 |
| AMPA-Pre,DCa | $1+x_1x_2+x_1^2+x_2^2$ | 0.106 |
| AMPA-Sch,DCa | $1+x_1+x_2$ | 0.147 |
| AMPA-Sch,DCa | $1+x_1x_2$ | 0.147 |
| AMPA-Sch,DCa | $1+x_1x_2+x_1^2+x_2^2$ | 0.15 |
| GABA,DCa | $1+x_1+x_2$ | 0.000108 |
| GABA,DCa | $1+x_1x_2$ | 0.00109 |
| GABA,DCa | $1+x_1x_2+x_1^2+x_2^2$ | 0.00112 |
| AMPA-Pre,DCb | $1+x_1+x_2$ | 0.082 |
| AMPA-Pre,DCb | $1+x_1x_2$ | 0.0821 |
| AMPA-Pre,DCb | $1+x_1x_2+x_1^2+x_2^2$ | 0.106 |
| AMPA-Sch,DCb | $1+x_1+x_2$ | 0.147 |
| AMPA-Sch,DCb | $1+x_1x_2$ | 0.147 |
| AMPA-Sch,DCb | $1+x_1x_2+x_1^2+x_2^2$ | 0.149 |
| GABA,DCb | $1+x_1+x_2$ | 0.00065 |
| GABA,DCb | $1+x_1x_2$ | 0.000651 |
| GABA,DCb | $1+x_1x_2+x_1^2+x_2^2$ | 0.000682 |
| DCa,DCb | $1+x_1+x_2$ | 0.000576 |
| DCa,DCb | $1+x_1x_2$ | 0.000578 |
| DCa,DCb | $1+x_1x_2+x_1^2+x_2^2$ | 0.00059 |
| AMPA-Pre | $1+x_1$ | 0.0815 |
| AMPA-Pre | $1+x_1+x_1^2$ | 0.106 |
| AMPA-Sch | $1+x_1$ | 0.147 |
| AMPA-Sch | $1+x_1+x_1^2$ | 0.149 |
| GABA | $1+x_1$ | 0.000581 |
| GABA | $1+x_1+x_1^2$ | 0.000605 |
| DCa | $1+x_1$ | 0.000506 |
| DCa | $1+x_1+x_1^2$ | 0.00051 |
| DCb | $1+x_1$ | $7.04 \cdot 10^{-5}$ |
| DCb | $1+x_1+x_1^2$ | $7.81 \cdot 10^{-5}$ |

**Table S5.**
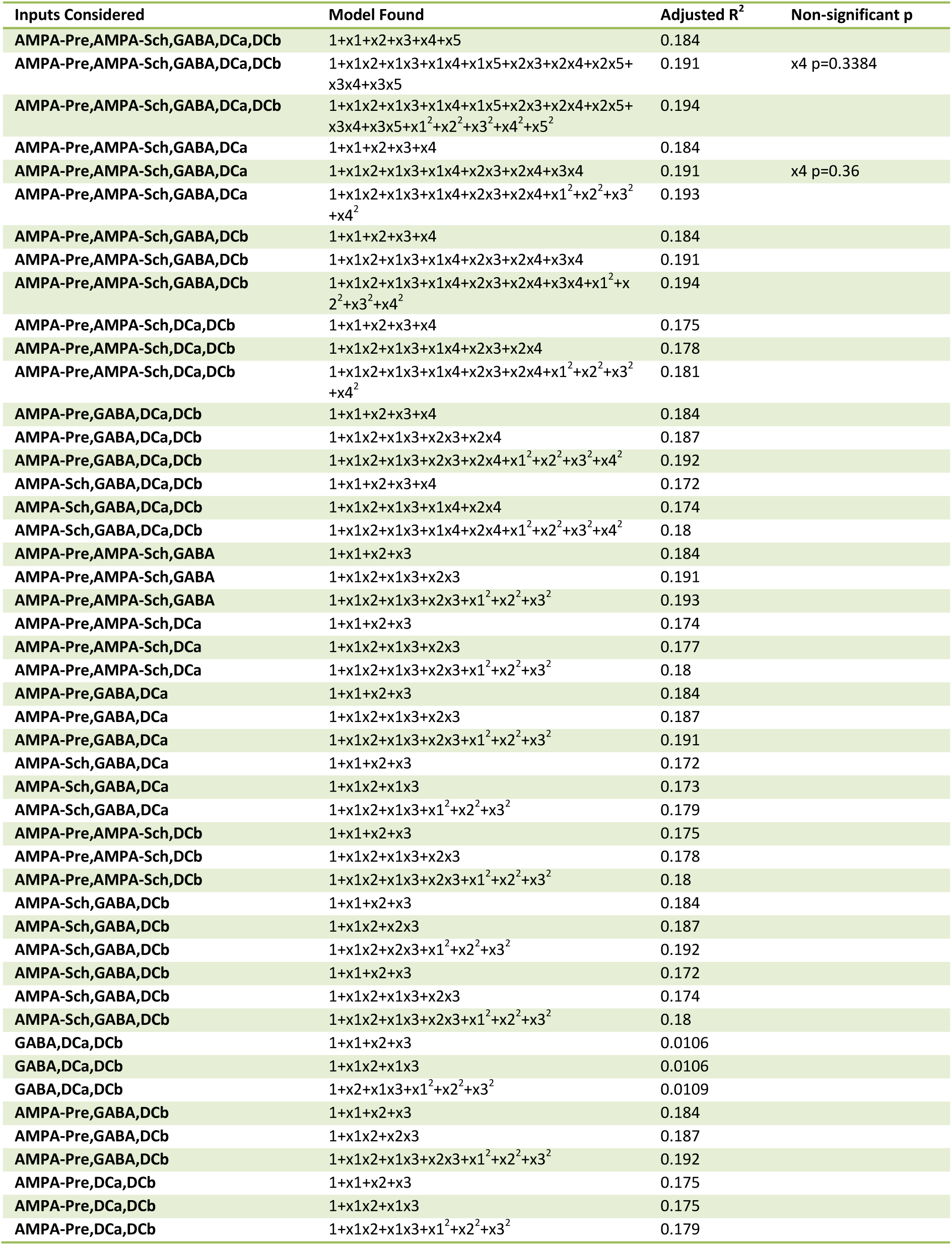
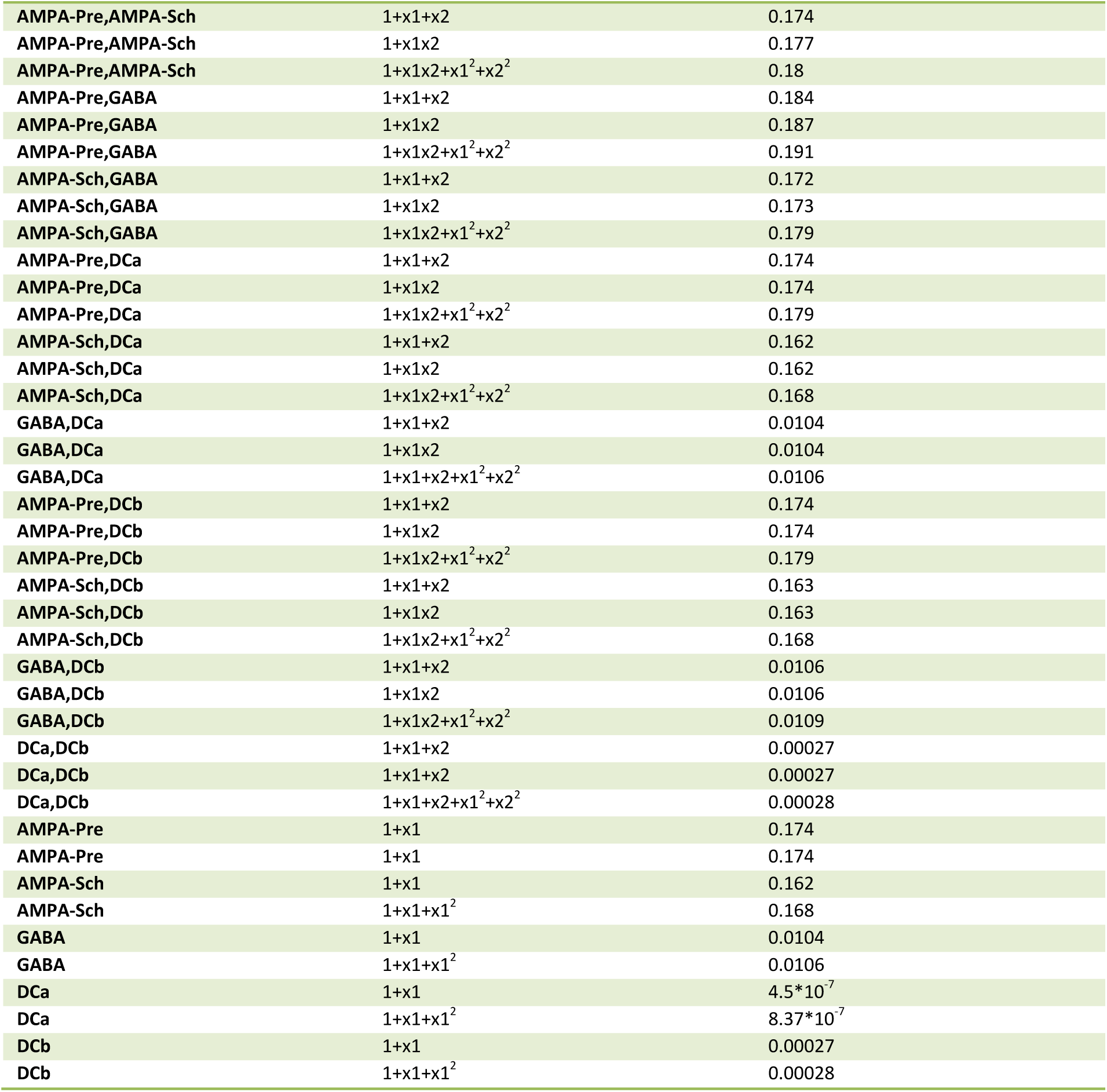
Linear regression models for R-activation of CA3-CA1 pyramidal cell pairs vs inputs. For each possible combination of the inputs considered (parameters of the model for the statistical linear fit), we found multiple fits to R-activation of CA3-CA1 pyramidal cell pairs. When only one input at a time was considered, we found linear models of linear terms and linear models of quadratic terms. When multiple inputs were considered, we found linear models for linear terms, interactions and quadratic terms. The models found are listed in the middle column, the value 1 represents the intercept at the origin, x1 represents the first input listed, x2 the second and so on. On occasion, parameters in the models did not pass the p-value significance requirement (p<0.05), in which case they were listed on the p-value last column. The adjusted R2 terms can be interpreted as a measure of how much of the variance in the data is captured by a given model. As can be seen from the values, there is a high noise component in the relationship between inputs and cell pair R-activation. However, the vast majority of models listed showed significant contributions of the inputs considered (by p-value<0.05 criterion).

| Inputs Considered | Model Found | Adjusted R<sup>2</sup> | Non-significant p |
| --- | --- | --- | --- |
| AMPA-Pre,AMPA-Sch,GABA,Dca,DCb | 1+x1+x2+x3+x4+x5 | 0.184 |  |
| AMPA-Pre,AMPA-Sch,GABA,Dca,DCb | 1+x1x2+x1x3+x1x4+x1x5+x2x3+x2x4+x2x5+x3x4+x3x5 | 0.191 | x4 p=0.3384 |
| AMPA-Pre,AMPA-Sch,GABA,Dca,DCb | 1+x1x2+x1x3+x1x4+x1x5+x2x3+x2x4+x2x5+x3x4+x3x5+x1<sup>2</sup>+x2<sup>2</sup>+x3<sup>2</sup>+x4<sup>2</sup>+x5<sup>2</sup> | 0.194 |  |
| AMPA-Pre,AMPA-Sch,GABA,Dca | 1+x1+x2+x3+x4 | 0.184 |  |
| AMPA-Pre,AMPA-Sch,GABA,Dca | 1+x1x2+x1x3+x1x4+x2x3+x2x4+x3x4 | 0.191 | x4 p=0.36 |
| AMPA-Pre,AMPA-Sch,GABA,Dca | 1+x1x2+x1x3+x1x4+x2x3+x2x4+x1<sup>2</sup>+x2<sup>2</sup>+x3<sup>2</sup>+x4<sup>2</sup> | 0.193 |  |
| AMPA-Pre,AMPA-Sch,GABA,DCb | 1+x1+x2+x3+x4 | 0.184 |  |
| AMPA-Pre,AMPA-Sch,GABA,DCb | 1+x1x2+x1x3+x1x4+x2x3+x2x4+x3x4 | 0.191 |  |
| AMPA-Pre,AMPA-Sch,GABA,DCb | 1+x1x2+x1x3+x1x4+x2x3+x2x4+x3x4+x1<sup>2</sup>+x2<sup>2</sup>+x3<sup>2</sup>+x4<sup>2</sup> | 0.194 |  |
| AMPA-Pre,AMPA-Sch,Dca,DCb | 1+x1+x2+x3+x4 | 0.175 |  |
| AMPA-Pre,AMPA-Sch,Dca,DCb | 1+x1x2+x1x3+x1x4+x2x3+x2x4 | 0.178 |  |
| AMPA-Pre,AMPA-Sch,Dca,DCb | 1+x1x2+x1x3+x1x4+x2x3+x2x4+x1<sup>2</sup>+x2<sup>2</sup>+x3<sup>2</sup>+x4<sup>2</sup> | 0.181 |  |
| AMPA-Pre,GABA,Dca,DCb | 1+x1+x2+x3+x4 | 0.184 |  |
| AMPA-Pre,GABA,Dca,DCb | 1+x1x2+x1x3+x2x3+x2x4 | 0.187 |  |
| AMPA-Pre,GABA,Dca,DCb | 1+x1x2+x1x3+x2x3+x2x4+x1<sup>2</sup>+x2<sup>2</sup>+x3<sup>2</sup>+x4<sup>2</sup> | 0.192 |  |
| AMPA-Sch,GABA,Dca,DCb | 1+x1+x2+x3+x4 | 0.172 |  |
| AMPA-Sch,GABA,Dca,DCb | 1+x1x2+x1x3+x1x4+x2x4 | 0.174 |  |
| AMPA-Sch,GABA,Dca,DCb | 1+x1x2+x1x3+x1x4+x2x4+x1<sup>2</sup>+x2<sup>2</sup>+x3<sup>2</sup>+x4<sup>2</sup> | 0.18 |  |
| AMPA-Pre,AMPA-Sch,GABA | 1+x1+x2+x3 | 0.184 |  |
| AMPA-Pre,AMPA-Sch,GABA | 1+x1x2+x1x3+x2x3 | 0.191 |  |
| AMPA-Pre,AMPA-Sch,GABA | 1+x1x2+x1x3+x2x3+x1<sup>2</sup>+x2<sup>2</sup>+x3<sup>2</sup> | 0.193 |  |
| AMPA-Pre,AMPA-Sch,Dca | 1+x1+x2+x3 | 0.174 |  |
| AMPA-Pre,AMPA-Sch,Dca | 1+x1x2+x1x3+x2x3 | 0.177 |  |
| AMPA-Pre,AMPA-Sch,Dca | 1+x1x2+x1x3+x2x3+x1<sup>2</sup>+x2<sup>2</sup>+x3<sup>2</sup> | 0.18 |  |
| AMPA-Pre,GABA,Dca | 1+x1+x2+x3 | 0.184 |  |
| AMPA-Pre,GABA,Dca | 1+x1x2+x1x3+x2x3 | 0.187 |  |
| AMPA-Pre,GABA,Dca | 1+x1x2+x1x3+x2x3+x1<sup>2</sup>+x2<sup>2</sup>+x3<sup>2</sup> | 0.191 |  |
| AMPA-Sch,GABA,Dca | 1+x1+x2+x3 | 0.172 |  |
| AMPA-Sch,GABA,Dca | 1+x1x2+x1x3 | 0.173 |  |
| AMPA-Sch,GABA,Dca | 1+x1x2+x1x3+x1<sup>2</sup>+x2<sup>2</sup>+x3<sup>2</sup> | 0.179 |  |
| AMPA-Pre,AMPA-Sch,DCb | 1+x1+x2+x3 | 0.175 |  |
| AMPA-Pre,AMPA-Sch,DCb | 1+x1x2+x1x3+x2x3 | 0.178 |  |
| AMPA-Pre,AMPA-Sch,DCb | 1+x1x2+x1x3+x2x3+x1<sup>2</sup>+x2<sup>2</sup>+x3<sup>2</sup> | 0.18 |  |
| AMPA-Sch,GABA,DCb | 1+x1+x2+x3 | 0.184 |  |
| AMPA-Sch,GABA,DCb | 1+x1x2+x2x3 | 0.187 |  |
| AMPA-Sch,GABA,DCb | 1+x1x2+x2x3+x1<sup>2</sup>+x2<sup>2</sup>+x3<sup>2</sup> | 0.192 |  |
| AMPA-Sch,GABA,DCb | 1+x1+x2+x3 | 0.172 |  |
| AMPA-Sch,GABA,DCb | 1+x1x2+x1x3+x2x3 | 0.174 |  |
| AMPA-Sch,GABA,DCb | 1+x1x2+x1x3+x2x3+x1<sup>2</sup>+x2<sup>2</sup>+x3<sup>2</sup> | 0.18 |  |
| GABA,Dca,DCb | 1+x1+x2+x3 | 0.0106 |  |
| GABA,Dca,DCb | 1+x1x2+x1x3 | 0.0106 |  |
| GABA,Dca,DCb | 1+x2+x1x3+x1<sup>2</sup>+x2<sup>2</sup>+x3<sup>2</sup> | 0.0109 |  |
| AMPA-Pre,GABA,DCb | 1+x1+x2+x3 | 0.184 |  |
| AMPA-Pre,GABA,DCb | 1+x1x2+x2x3 | 0.187 |  |
| AMPA-Pre,GABA,DCb | 1+x1x2+x1x3+x2x3+x1<sup>2</sup>+x2<sup>2</sup>+x3<sup>2</sup> | 0.192 |  |
| AMPA-Pre,Dca,DCb | 1+x1+x2+x3 | 0.175 |  |
| AMPA-Pre,Dca,DCb | 1+x1x2+x1x3 | 0.175 |  |
| AMPA-Pre,Dca,DCb | 1+x1x2+x1x3+x1<sup>2</sup>+x2<sup>2</sup>+x3<sup>2</sup> | 0.179 |  |

|  |  |  |
| --- | --- | --- |
| AMPA-Pre,AMPA-Sch | 1+x1+x2 | 0.174 |
| AMPA-Pre,AMPA-Sch | 1+x1x2 | 0.177 |
| AMPA-Pre,AMPA-Sch | 1+x1x2+x1<sup>2</sup>+x2<sup>2</sup> | 0.18 |
| AMPA-Pre,GABA | 1+x1+x2 | 0.184 |
| AMPA-Pre,GABA | 1+x1x2 | 0.187 |
| AMPA-Pre,GABA | 1+x1x2+x1<sup>2</sup>+x2<sup>2</sup> | 0.191 |
| AMPA-Sch,GABA | 1+x1+x2 | 0.172 |
| AMPA-Sch,GABA | 1+x1x2 | 0.173 |
| AMPA-Sch,GABA | 1+x1x2+x1<sup>2</sup>+x2<sup>2</sup> | 0.179 |
| AMPA-Pre,DCa | 1+x1+x2 | 0.174 |
| AMPA-Pre,DCa | 1+x1x2 | 0.174 |
| AMPA-Pre,DCa | 1+x1x2+x1<sup>2</sup>+x2<sup>2</sup> | 0.179 |
| AMPA-Sch,DCa | 1+x1+x2 | 0.162 |
| AMPA-Sch,DCa | 1+x1x2 | 0.162 |
| AMPA-Sch,DCa | 1+x1x2+x1<sup>2</sup>+x2<sup>2</sup> | 0.168 |
| GABA,DCa | 1+x1+x2 | 0.0104 |
| GABA,DCa | 1+x1x2 | 0.0104 |
| GABA,DCa | 1+x1+x2+x1<sup>2</sup>+x2<sup>2</sup> | 0.0106 |
| AMPA-Pre,DCb | 1+x1+x2 | 0.174 |
| AMPA-Pre,DCb | 1+x1x2 | 0.174 |
| AMPA-Pre,DCb | 1+x1x2+x1<sup>2</sup>+x2<sup>2</sup> | 0.179 |
| AMPA-Sch,DCb | 1+x1+x2 | 0.163 |
| AMPA-Sch,DCb | 1+x1x2 | 0.163 |
| AMPA-Sch,DCb | 1+x1x2+x1<sup>2</sup>+x2<sup>2</sup> | 0.168 |
| GABA,DCb | 1+x1+x2 | 0.0106 |
| GABA,DCb | 1+x1x2 | 0.0106 |
| GABA,DCb | 1+x1x2+x1<sup>2</sup>+x2<sup>2</sup> | 0.0109 |
| DCa,DCb | 1+x1+x2 | 0.00027 |
| DCa,DCb | 1+x1+x2 | 0.00027 |
| DCa,DCb | 1+x1+x2+x1<sup>2</sup>+x2<sup>2</sup> | 0.00028 |
| AMPA-Pre | 1+x1 | 0.174 |
| AMPA-Pre | 1+x1 | 0.174 |
| AMPA-Sch | 1+x1 | 0.162 |
| AMPA-Sch | 1+x1+x1<sup>2</sup> | 0.168 |
| GABA | 1+x1 | 0.0104 |
| GABA | 1+x1+x1<sup>2</sup> | 0.0106 |
| DCa | 1+x1 | 4.5*10<sup>-7</sup> |
| DCa | 1+x1+x1<sup>2</sup> | 8.37*10<sup>-7</sup> |
| DCb | 1+x1 | 0.00027 |
| DCb | 1+x1+x1<sup>2</sup> | 0.00028 |

